# The plant immune receptor LORE binds agonistic and antagonistic 3-hydroxy fatty acid ligands via a dynamic loop in its G-type lectin domain

**DOI:** 10.64898/2026.06.08.730953

**Authors:** Lin-Jie Shu, Alessandro Nicoli, Fan-Yu Yu, Olga O. A. Thiry, Gabriel Deslandes-Hérold, Tom Luethi, Antonella Di Pizio, Stefanie Ranf

## Abstract

The *Arabidopsis thaliana* S-domain receptor kinase LORE senses bacterial medium-chain 3-hydroxy fatty acids (mc-3-OH-FAs) as microbe-associated molecular patterns to activate pattern-triggered immunity. How LORE recognises these fatty acid ligands at the molecular level remains unknown. Here, we combined protein structure prediction, protein–ligand interaction modelling and molecular dynamics (MD) simulations with ligand-binding assays using chimeric and mutant receptor ectodomains, and functional analysis of receptor activation to characterise the mc-3-OH-FA binding mechanism. Domain-swap experiments between LORE and its non-binding paralog *At*SD1-23 identify the lectin 2 (L2) domain as the ligand-binding domain. Mutational analysis and reverse engineering confirm a hydrophobic pocket in the L2 core as the primary ligand-binding site. Multiple walker Supervised MD (mwSuMD) simulations reveal that the acyl tail enters the pocket first, whilst polar interactions between the headgroup and a flexible L2 loop guide and stabilise the bound state. In support of this model, 3-OH-C10:0 analogues with bulky headgroup modifications dock into the pocket but act as antagonists, presumably by preventing the loop from adopting the conformation required for signalling. Together, these data suggest that the flexible L2 loop has multiple functions: it acts as a dynamic gate regulating pocket access, provides essential anchoring points once the ligand is bound, and contributes to receptor activation. These findings provide a mechanistic framework for immunogenic mc-3-OH-FA sensing by LORE.

## INTRODUCTION

Plants employ a multilayered innate immune system to restrict microbial invasion. A critical component of this system operates at the cell surface, where pattern recognition receptors (PRRs) sense conserved microbe-associated molecular patterns (MAMPs) and initiate pattern-triggered immunity (PTI). PTI acts in concert with intracellular effector-triggered immunity (ETI) to form an integrated immune network (Ngou *et al*., 2021; Yuan *et al*., 2021). PRRs are typically receptor-like kinases (RLKs) or receptor-like proteins (RLPs) and are classified according to the architecture of their extracellular domain (ECD) into, for example, leucine-rich repeat (LRR), Lysin Motif (LysM), and S-domain (SD) receptors (Shiu & Bleecker, 2001a; Dievart *et al*., 2020). RLKs comprise an ECD, a single-span transmembrane domain (TMD), and an intracellular domain (ICD) with a serine/threonine/tyrosine kinase domain, whereas RLPs lack a major intracellular signalling module. Since PTI activation is broadly conserved across plant species, PRRs are prime candidates for engineering durable, broad-spectrum disease resistance in crop plants (Boutrot & Zipfel, 2017). Realising this potential, however, requires a comprehensive mechanistic understanding of how individual PRR classes perceive their ligands and transduce the signal across the membrane.

Among plant RLK families, LRR-RLKs have been most intensively characterised at the structural and mechanistic level (Hohmann *et al*., 2017; Ngou *et al*., 2026). Structural and biochemical studies have revealed different receptor activation mechanisms within this class. Many LRR-RLKs are activated through ligand-induced recruitment of SOMATIC EMBRYOGENESIS RECEPTOR-LIKE KINASE (SERK) family co-receptors: in some cases, the ligand directly bridges the receptor–co-receptor interface, acting as a ‘molecular glue’ that stabilises the complex (e.g. FLS2/flg22/BAK1, HAE/IDA/BAK1); in others, such as BRI1, the ligand first binds to a dedicated island domain within the LRR ECD and subsequently promotes co-receptor association (Fan *et al*., 2016; Song *et al*., 2017; Hohmann *et al*., 2018). In contrast, SD-RLKs, also termed bulb-type or G-type lectin RLKs, although constituting the second-largest RLK family with approximately 40 members in *Arabidopsis thaliana*, 70 in tomato, 100 in rice, and 120 in soybean (Shiu & Bleecker, 2001b; Shiu *et al*., 2004; Vaid *et al*., 2012; Teixeira *et al*., 2018; Dievart *et al*., 2020), remain largely uncharacterised at the mechanistic level. Their ECDs are defined by tandem *Galanthus nivalis* agglutinin (G)-type lectin-like domains (L1 and L2), an epidermal growth factor-like domain (EGF), and a plasminogen-apple-nematode domain (PAN) (Shiu & Bleecker, 2003; Vaid *et al*., 2012; Ma *et al*., 2016; Teixeira *et al*., 2018; Dievart *et al*., 2020). The family was initially defined through the S-LOCUS RECEPTOR KINASE (SRK) of *Brassica oleracea*, which mediates haplotype-specific recognition of pollen-expressed SCR peptides in the self-incompatibility response (Nasrallah *et al*., 1987; Stein *et al*., 1991; Naithani *et al*., 2007; Ivanov *et al*., 2010). Beyond reproduction, SD-RLKs have since been implicated in a diverse array of biological processes, including abiotic stress responses, tissue development, pathogen defence, and symbiosis across multiple plant species (Charpenteau *et al*., 2004; Chen *et al*., 2006; Bonaventure, 2011; Chen *et al*., 2013; Cheng *et al*., 2013; Sun *et al*., 2013; Zhao *et al*., 2013; Catanzariti *et al*., 2015; Ranf *et al*., 2015; Zou *et al*., 2015; Fan *et al*., 2018; Sun *et al*., 2018; Labbé *et al*., 2019; Pan *et al*., 2020; Kato *et al*., 2022; Pi *et al*., 2022; Zhang *et al*., 2022; Bao *et al*., 2023; Guo *et al*., 2023; Liu *et al*., 2023; Micol-Ponce *et al*., 2023; Xu *et al*., 2023; Zhou *et al*., 2023; Chien *et al*., 2024; Li *et al*., 2024; Chen *et al*., 2025; Niu *et al*., 2025; Choi *et al*., 2026; Li *et al*., 2026; Shen *et al*., 2026). They form a rapidly evolving class, with individual subclades expanding largely through tandem gene duplication (Xing *et al*., 2013). Despite their functional diversity and biological importance, how SD-RLKs perceive their ligands at the molecular level remains largely unknown.

We have previously identified the Brassicaceae-specific SD-RLK LIPOOLIGOSACCHARIDE-SPECIFIC REDUCED ELICITATION (LORE; alias SD1-29) as a PRR that senses medium-chain 3-hydroxy fatty acids (mc-3-OH-FAs) and their derivatives 3-(3-hydroxyalkanoyloxy)alkanoic acids (HAAs) as MAMPs in antibacterial immunity (Ranf *et al*., 2015; Kutschera *et al*., 2019; Schellenberger *et al*., 2021; Shu *et al*., 2026a). mc-3-OH-FAs with aliphatic tails of 8 to 13 carbons, most potently 3-hydroxydecanoic acid (3-OH-C10:0), trigger LORE-dependent immune responses in Brassicaceae, including cytoplasmic calcium ([Ca^2+^]_cyt_) signalling, reactive oxygen species (ROS) production, activation of mitogen-activated protein kinases (MAPKs), and defence gene expression (Kutschera *et al*., 2019; Schellenberger *et al*., 2021; Shu *et al*., 2026a). By contrast, long-chain 3-OH-FAs with aliphatic tails of 14 carbons or longer are inactive. This chain-length discrimination is an intrinsic property of the LORE ECD, as this specificity is retained when LORE is transferred to solanaceous species that otherwise lack the ability to sense mc-3-OH-FAs (Ranf *et al*., 2015; Kutschera *et al*., 2019; Eschrig *et al*., 2024a; Eschrig *et al*., 2024b; Shu *et al*., 2026a). A detailed structure-activity analysis further revealed that modifications to the carboxylate headgroup, the position of the hydroxyl group, and the absolute configuration all influence elicitor activity (Kutschera *et al*., 2019). mc-3-OH-FAs are found in relevant quantities in plant-associated Proteobacteria, particularly β- and γ-Proteobacteria, where they serve as biosynthetic precursors for lipopolysaccharides and other cellular components; however, the precise mechanisms by which they are released into the apoplast and whether the free form predominates *in planta* remain unknown (Kutschera *et al*., 2019; Shu *et al*., 2026a). mc-3-OH-FA sensing is specific to LORE and is not shared by closely related SD-RLK paralogs. *At*SD1-23, the closest paralog of *At*LORE with high ECD sequence similarity, neither binds 3-OH-C10:0 nor responds to mc-3-OH-FA elicitation (Ranf *et al*., 2015; Shu *et al*., 2021; Shu *et al*., 2026a). LORE-mediated mc-3-OH-FA sensing is conserved across Brassicaceae, and the chain-length preference of LORE orthologs correlates with the mc-3-OH-FA profiles of plant-associated Proteobacteria (Shu *et al*., 2026a). Together with the identification of the Arabidopsis SD-RLK RDA2/SD1-12 as a PRR for sphingolipid-derived 9-methyl sphingoid base (Kato *et al*., 2022), the functional characterisation of LORE supports an emerging role for SD-RLKs as sensors of lipidic immunogenic metabolites. As one of a few SD-RLKs with a known activating ligand, LORE represents an important prototype for this large and underexplored receptor family.

Mechanistically, SRKs are the best studied SD-RLKs. Crystal structures of the SRK ECD in complex with its cognate SCR/SP11 ligand have been determined for two *Brassica rapa* haplotypes (Ma *et al*., 2016; Murase *et al*., 2020), revealing that SCR recognition is mediated through hypervariable regions of the L2 and EGF-like domains and that ligand binding stabilises receptor homodimerisation into a 2:2 heterotetrameric complex. LORE likewise requires homodimerisation through its ECD and TMD, combined with a catalytically active kinase domain, for receptor activation (Luo *et al*., 2020; Wang *et al*., 2023; Eschrig *et al*., 2024b). Notably, LORE orthologs from *A. lyrata* and *A. halleri* have 3-OH-C10:0-binding capacity but fail to homodimerise and are consequently non-functional in immune signalling, illustrating that ligand binding and receptor activation can be uncoupled (Eschrig *et al*., 2024b; Shu *et al*., 2026a). The SCR/SP11 ligands of SRK are small cysteine-rich peptides (∼50 amino acids) with a defensin-like fold, which are chemically and structurally distinct from mc-3-OH-FAs (Ma *et al*., 2016; Murase *et al*., 2020). The SRK–SCR interaction thus provides no direct template for understanding how LORE recognises its small lipidic ligand or how ligand binding leads to receptor homomerisation and activation. Here, we combined AI- and homology-based structural modelling of the LORE-ECD with molecular docking, multiple walker Supervised Molecular Dynamics (mwSuMD), and targeted mutagenesis to define the mc-3-OH-FA binding site and characterise the binding mechanism. Our results identify a hydrophobic pocket within the L2 domain as the primary 3-OH-FA binding site, reveal a binding mechanism in which the flexible L2 loop shapes the binding site and stabilises the bound state, and demonstrate that 3-OH-C10:0 analogues with bulky headgroup modifications act as antagonists of LORE signalling. Together, these findings provide a mechanistic framework for lipidic ligand recognition by an SD-RLK and a structural basis for understanding how ligand binding may drive the receptor conformational changes required for activation.

## RESULTS

### 3-OH-FA ligands bind to a central cavity in the lectin 2 domain of LORE

Since no experimental structure for the LORE ECD has been determined to date, we generated three-dimensional models using both homology-based and AI-based approaches. Homology models were built with MODELLER (Šali & Blundell, 1993) using the crystal structures of SRK8 (PDB: 6KYW) and SRK9 (PDB: 5GYY) as templates (**Fig. S1A, B**). An independent structural prediction was performed using AlphaFold2 (AF2) (Jumper *et al*., 2021) (**Fig. S1C**). A comparative analysis of the resulting models revealed high conservation of the global architecture, with overall Root Mean Square Deviation (RMSD) of the backbone Cα values ranging from 1.8 to 3.2 Å (**Fig. 1A; Fig. S2A**). This level of agreement across different modelling platforms indicates a consistent and reliable protein fold. Differences identified in loop regions may reflect the inherent flexibility of these loops or differences in template-specific conformations (**Fig. S2B**). Pocket detection identified a primary binding cavity located within the L2 region (**Fig. 1B; S3A**). This cavity consistently ranked as the optimal binding site and exhibited the highest druggability across all models (**Fig. 1B**), indicating that it is the most likely site for ligand interaction. To functionally validate the predicted 3-OH-FA binding domain in *At*LORE, we tested chimeric ECDs of *At*LORE and the closely related non-3-OH-FA-binding *At*SD1-23 with swapped subdomains for ligand binding. We expressed the ECDs of *At*LORE, *At*SD1-23 and various chimera as apoplastic mCherry fusion proteins in *N. benthamiana* (**Fig. S4A, B**) and assessed 3-OH-C10:0 binding using a ligand-depletion assay (**Fig. 1C**) (Shu *et al*., 2021). Not all combinations could be successfully expressed, suggesting that certain combinations of *At*LORE and *At*SD1-23 subdomains are incompatible. Of the expressed ones, only chimeric ECDs comprising the complete L2 domain of *At*LORE showed 3-OH-C10:0-binding activity, while those with a partial or complete L2 domain from *At*SD1-23 did not. Receptor activity of the *At*LORE/*At*SD1-23 chimeric ECDs fused to *At*LORE-TMD-ICD was tested upon transient expression in *N. benthamiana* by measuring ROS production in response to elicitation with 3-OH-C10:0 (**Fig. S4C**). In agreement with the binding analyses, only the chimera comprising the complete L2 domain of *At*LORE showed significant 3-OH-C10:0-induced signalling activity, while activity was strongly reduced or abolished in chimera comprising a partial or complete L2 domain from *At*SD1-23, respectively. These findings identify the LORE L2 domain as the 3-OH-C10:0-binding module and support the predicted pocket as the 3-OH-C10:0 binding site.

**Figure 1.**
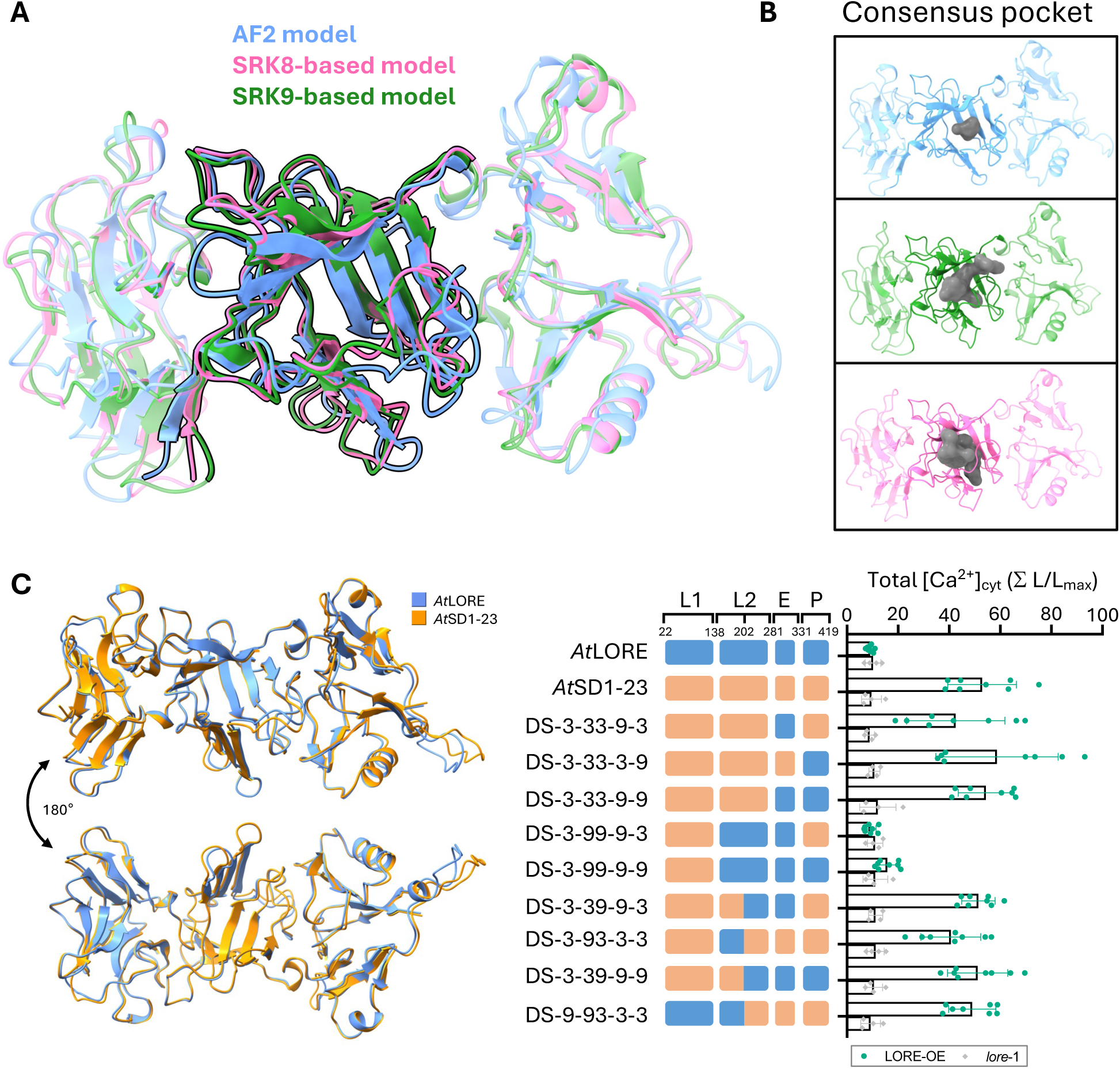
Mapping of the 3-OH-FA-binding domain in the *At*LORE ectodomain. **A** Superimposition of three predicted models of the LORE ECD: AlphaFold2 (AF2) predicted model (blue), SRK8-based homology model (pink), and SRK9-based homology model (green). **B** Individual representations of each model illustrating the location of the consensus binding pocket (grey surface) identified across all three models using three different binding pocket detection methods. AF2 model: Fpocket (volume = 189.20 Å3, score = 0.297, druggability = 0.516/1) and Sitemap (volume = 53, score = 1.882, druggability = 1.14). SRK8-based model: Fpocket (volume = 1028.53 Å3, score = 0.499, druggability = 0.985/1) and Sitemap (volume = 229, score = 1.232, druggability = 1.232). SRK9-based: Fpocket (volume = 1223.98 Å3, score = 0.208, druggability = 0.998/1) and Sitemap (volume = 341.19, score = 1.268, druggability = 1.264). **C** Left panel: superimposition of AF2 models of the ECDs of *At*LORE (AF-O64782-F1-v2, 22-417, blue) and *At*SD1-23 (AF-O64781-2-F1-v2, 42-438, orange). Right panel: Result of ligand-depletion assay using the indicated ECDs or subdomain chimera performed with 500 nM 3-OH-C10:0 and concentrated apoplastic washing fluids (AWFs) from *N. benthamiana* containing the indicated ECD-mCherry fusion proteins (1.5 mg/mL total protein concentration; anti-RFP immunoblots shown in **Figure S4**). Increases in [Ca2+]_cyt_ were measured in LORE overexpressing (LORE-OE/*lore*-1) and knockout (*lore-*1) Arabidopsis seedlings treated with filtrates from the depletion assay (mean ± SD with individual data points; pooled data from two independent experiments; total number of seedlings used per ECD: LORE-OE: n=8, *lore*-1: n=4). DS; domain swap; L1, lectin 1; L2a/b, lectin 2a/b; E, epidermal growth factor (EGF)-like; P, plasminogen-apple-nematode (PAN); 3 indicates subdomains of *At*SD1-23, visualised as orange boxes; 9 indicates subdomains of *AtLORE,* visualised as blue boxes; numbers above boxes indicate domain swap sites according to *At*LORE numbering.

### Mutation of key residues in the binding pocket abolishes ligand binding and receptor activity

To identify residues responsible for ligand sensing, we combined sequence-based and structure-based analyses. We have previously shown that 3-OH-C10:0 binding is specific to LORE and highly conserved among LORE orthologs across the Brassicaceae family (Shu *et al*., 2026a). A multiple sequence alignment (MSA) of 3-OH-C10:0-binding LORE and non-binding SD1-23 orthologs revealed several single amino acid polymorphisms (SAPs) that distinguish LORE and SD1-23 orthologs (**Fig. 2A; S5A**). Such SAPs were enriched specifically in the L2 domain compared with other ECD subdomains. The MSA also identified various residues that were highly conserved among all LORE and SD1-23 orthologs (**Fig. 2A**). Candidate residues were mapped onto the predicted binding pockets of the *At*LORE models to assess their orientation and potential for ligand contact (**Fig. 2B, C; S5B, C**). This analysis distinguished between residues oriented towards the interior of the binding cavity and those pointing outwards. Filtering based on orientation and chemical properties identified five polar residues within the pocket: T190, S221, S224, Y241, and S250 (**Fig. 2B, C**). These residues were prioritised as potential “anchoring points” that could form key hydrogen bonds or other polar interactions with the ligand. However, S221 and S224 are in the flexible L2 loop. Their orientation varied across the models in the absence of the ligand, and they were not oriented toward the pocket in the AF2 model (**Fig. S5B**). In the non-binders, position 190 is occupied by proline; in the binders, it is leucine when threonine is at position 204 (**Fig. 2C; S5A**). Beyond the polar anchors, I184 was identified as being in a key position at the extremity of the pocket (**Fig. 2B, C**), suggesting that it plays an important role in determining the shape of the pocket. Interestingly, near this site, LORE has a conserved G237 residue, whereas non-binding orthologs have an alanine at this position (**Fig. 2A-C; S5A**), which would further constrict the pocket. To functionally assess the role of selected candidate residues, we tested single and higher-order mutants for 3-OH-C10:0 binding and 3-OH-C10:0-induced signalling. For residues that differ among *At*LORE, LORE orthologs, and *At*SD1-23, we selected mutations matching those of *At*SD1-23 (I184F, G237A, T190P) or LORE orthologs (T190L, C204T) (**Fig. 2A; S5A**), while conserved residues were mutated to alanine (S224A, Y241A). Ligand binding to mutated *At*LORE-ECDs, expressed as apoplastic mCherry fusion proteins in *N. benthamiana,* was tested using the ligand-depletion assay (**Fig. 2D**). All mutant variants were expressed and secreted to the *N. benthamiana* apoplast (**Fig. S6**). T190P, T190L, T190L-C204T or S250A mutants bind 3-OH-C10:0, thus excluding two possible subpockets in the L2 cavity (**Fig. S3B**). 3-OH-C10:0 binding of Y241A and I184F is reduced, whereas the combination of I184F and G237A, like in *At*SD1-23, abolishes 3-OH-C10:0 binding to the *At*LORE-ECD (**Fig. 2D**). This confirms not only the ligand-binding pocket in the L2 domain but also identifies key residues that define its boundaries. Next, we tested the receptor activity of the *At*LORE mutants upon transient expression in *N. benthamiana* by measuring ROS production in response to elicitation with 3-OH-C10:0 (**Fig. 2E**). In agreement with the 3-OH-C10:0 binding analyses, I184F reduces receptor activity, while additional mutation of G237A abolishes activity. L154V and L274F, two other SAPs of SD1-23 relative to LORE, located at the pocket extremity and the opening, respectively, also reduce receptor activity, whereas L276A does not (**Fig. 2A, B, E; S5A**). S250A, T190L, and T190L-C204T do not affect receptor activity. Interestingly, T190P, although not impaired in binding 3-OH-C10:0, reduces 3-OH-C10:0-induced LORE activity (**Fig. 2D, E**). Similarly, Y241A, despite still showing partial 3-OH-C10:0 binding capacity, abolishes receptor activity (**Fig. 2D, E**). S224A, located in the flexible L2 loop, also abolishes receptor activity (**Fig. 2E**). In conclusion, our analyses identify the L2 domain as the functional ligand-binding pocket of *At*LORE, where residues I184 and G237 act as key determinants of pocket shape and binding specificity. The results reveal a critical distinction between ligand recognition and receptor activation, highlighting residues like Y241 as essential for signal transduction. Ultimately, these findings provide a molecular map of how specific SAPs govern the evolutionary divergence of mc-3-OH-FA sensing across the Brassicaceae family.

**Figure 2.**
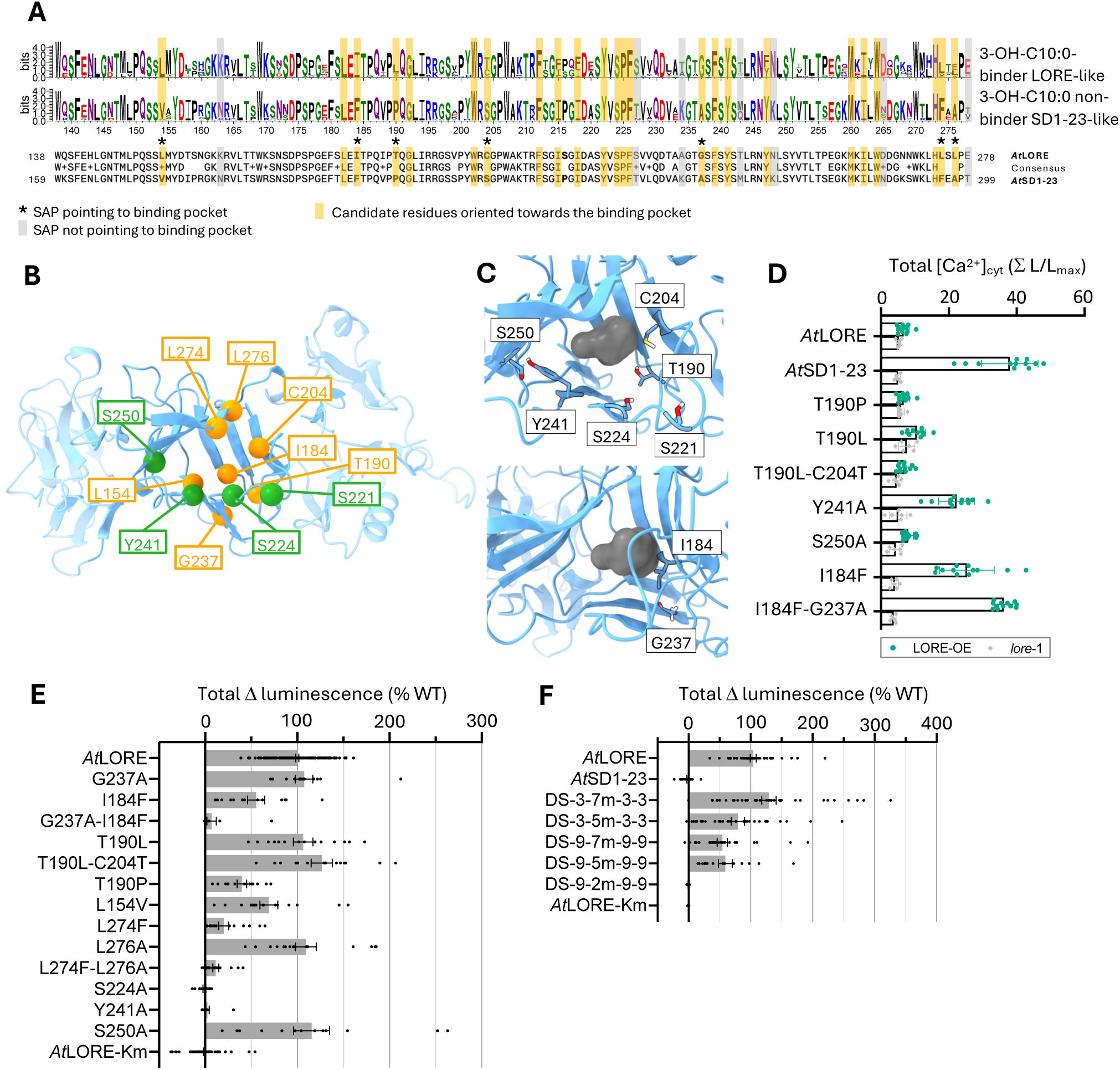
Identification of key residues in the lectin 2 domain contributing to 3-OH-FA binding. **A** Upper panel: WebLogos of multiple sequence alignments of the L2 domain of LORE or SD1-23 orthologs (numbering according to *At*LORE). Lower panel: sequence alignment of the L2 domain of *At*LORE and *At*SD1-23. **B** AF2 model of *At*LORE (AF-O64782-F1-v2, 22-417, blue) with the L2 domain highlighted in blue, SAPs relative to *At*SD1-23 oriented towards the binding pocket in L2 are depicted in orange and polar residues in green. **C** Zoom-in view of the consensus binding pocket in the AF2 model. Selected residue from the identified L2 cavity are depicted as blue sticks. **D** Result of ligand-depletion assay using ECDs of *At*LORE, *At*SD1-23 or different *At*LORE mutants performed with 500 nM 3-OH-C10:0 and concentrated apoplastic washing fluids (AWFs) from *N. benthamiana* containing the indicated ECD-mCherry fusion proteins (1.5 mg/mL total protein concentration; anti-RFP immunoblots shown in **Figure S6**). Increases in [Ca2+]_cyt_ were measured in LORE overexpressing (LORE-OE/*lore*-1) and knockout (*lore-*1) Arabidopsis seedlings treated with filtrates from the depletion assay. Graph shows pooled data from three independent experiments presented as mean ± SD with individual data points; total number of seedlings used per ECD: LORE-OE: n=12, *lore*-1: n=6. **E, F** ROS production was measured 2 dpi in leaf disks from *N. benthamiana* leaves expressing *At*LORE or the indicated *At*LORE mutants (**E**) or chimera (**F**) after elicitation with 5 µM 3-OH-C10:0 or water as control. Four mutants and *At*LORE were expressed on the same leaves. Kinase-mutated *At*LORE-Km was used as a negative control. Graph shows pooled data from two independent experiments, expressed as % of wild-type response and presented as mean ± SEM with individual data points, from n ≥ 6 leaf disks per construct across a total of 3-4 replicate leaves from 2 plants. DS; domain swap; L1, lectin 1; L2, lectin 2; E, epidermal growth factor-like; P, plasminogen-apple-nematode; SAP, single amino acid polymorphism; 3 indicates subdomains of *At*SD1-23; 9 indicates subdomains of *At*LORE; 2m/5m/7m indicates *At*SD1-23 L2 domain with 2, 5, or 7 SAPs converted to the respective *At*LORE residues, respectively.

### Reverse L2 engineering converts SD1-23 into a functional mc-3-OH-FA receptor

Several residues of the L2 domain that affect 3-OH-C10:0 binding and/or receptor activation were identified from SAPs in the non-binding *At*SD1-23 relative to *At*LORE, while other residues are conserved among both proteins (**Fig. 2A**). We hypothesised that changing residues critical to 3-OH-C10:0 binding in the L2 domain of *At*SD1-23 to the respective *At*LORE residues should convert it into a 3-OH-C10:0 binding domain. Considering potential incompatibilities among other subdomains of *At*SD1-23 and *At*LORE, we created different chimeric ECDs all sharing a converted *At*SD1-23 L2 domain, where all seven SAPs located in the binding pocket are changed to the respective *At*LORE residues (**Fig. 2A, B**; L2^7m^: V175L, F205I, P211T, S225C, A258G, F295L, and A297L), the five SAPs that reduce activity of *At*LORE (**Fig. 2E**; L2^5m^: V175L, F205I, P211T, A258G, and F295L), or only the two SAPs that in combination abolish ligand binding (**Fig. 2D**; L2^2m^: F205I, A258G). Receptor activity of these chimeric ECDs fused to *At*LORE-TMD-ICD was tested upon transient expression in *N. benthamiana* by measuring ROS production in response to elicitation with 3-OH-C10:0. All chimera with a converted L2^7m^ or L2^5m^ domain gained 3-OH-C10:0 receptor function, while those with only F205I-A258G (L2^2m^) did not (**Fig. 2F**). The results provide direct experimental support for the predicted binding pocket and confirm that the L2 domain is both necessary and sufficient to encode mc-3-OH-FA sensing specificity. Furthermore, since L2^7m^ engineering converts the *At*SD1-23 ECD into a fully functional 3-OH-C10:0 receptor domain, its remaining subdomains must be sufficiently conserved to enable receptor activation and signalling.

### 3-OH-C10:0 binding to LORE involves the flexible L2 loop

To characterise the binding process and sample the conformational landscape of the *At*LORE-ECD:3-OH-C10:0 complex, we employed a three-pronged computational strategy to extensively sample binding site side-chain flexibility: (1) Induced-Fit Docking (IFD) (**Fig. S7**) (Sherman *et al*., 2006a; Sherman *et al*., 2006b), (2) Boltz-2 cofolding (**Fig. S8**) (Passaro *et al*., 2025), and (3) mwSuMD (**Fig. S9**) (Deganutti *et al*., 2025). IFD samples ligand binding poses by optimising protein-ligand complementarity within a defined pocket, capturing low-energy conformations while allowing protein residue flexibility. Boltz-2 uses protein sequence and ligand structure information to predict the bound complex, providing a structure that is independent of the experimental pocket definition and capable of placing the ligand without geometric constraints. Instead, mwSuMD was applied to simulate the binding pathway from the solvent to the pocket. Here, the ligand reaches the pocket spontaneously, supervised only by a collective variable that rewards proximity to the protein, without any prior assumption on the bound conformation.

A comparative analysis of IFD, Boltz-2 and mwSuMD results was performed to identify and characterise a consensus binding mode. In two out of three mwSuMD replicas (R1 and R3), the ligand successfully entered the predicted binding pocket (**Fig. S9**). In both cases, the hydrophobic tail of 3-OH-C10:0 enters the cavity first and adopts a stable, uniform orientation within the hydrophobic core, while the polar headgroup displays different conformations (**Fig. 3A**). We analysed the datasets of structures obtained from Boltz-2 and IFD, using the final bound states from the mwSuMD simulations as references. Across its heterogeneous sampling, IFD successfully samples poses similar (RMSD < 3 Å) to mwSuMD R1 (**Fig. 3B; S7**). Impressively, the extensive Boltz-2 co-folding sampling of 3-OH-C10:0 and the LORE ECD (1250 poses) converged very closely on a binding pose that is consistent with the mwSuMD R1 last productive snapshot (**Fig. 3B; S8**). This provided independent, sequence-based support for the predicted binding mode, thereby reinforcing the structural conclusions drawn from the mutational analysis.

**Figure 3.**
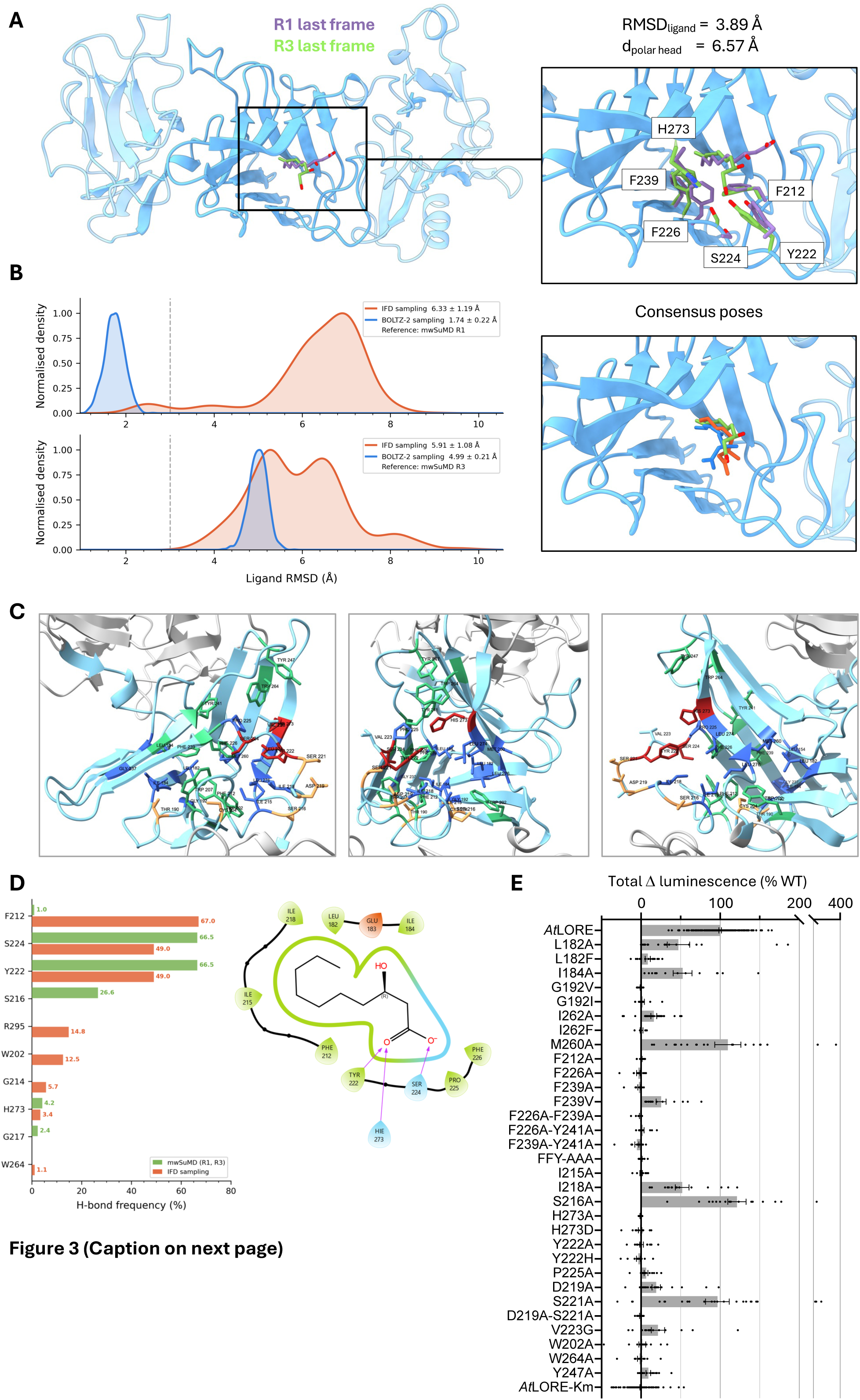
Characterisation of the 3-OH-C10:0 binding process to the LORE ectodomain. **A** End point of the two productive mwSuMD simulations (R1 and R3). Protein residues are depicted as blue cartoons, ligand and protein side chains as green (R1) and purple (R3) sticks. **B** Left panel: ligand heavy atoms RMSD of generated poses (IFD in orange, Boltz in blue). The RMSDs are shown as a distribution, using the positions of R1 and R3 as references (dashed line: RMSD cut off = 3 Å). Right panel: consensus poses between mwSuMD, IFD and Boltz-2. **C** Candidate residues forming the binding cavity and the L2 loop, visualised in the AF2 model of LORE. **D** Left panel: hydrogen bond analysis plot of binding site residues from IFD and mwSuMD. Right panel: 2D ligand protein interactions diagram from Boltz-2 best pose (RMSD to R1 = 1.13 Å). **E** ROS production was measured 2 dpi in leaf disks from *N. benthamiana* expressing *At*LORE or mutants elicited with 5 µM 3-OH-C10:0 or water as control. Four mutants and *At*LORE were expressed on the same leaves with kinase-mutated *At*LORE-Km as a negative control. Graph shows pooled data from two independent experiments, expressed as % of wild-type response and presented as mean ± SEM with individual data points, from n ≥ 6 leaf disks per construct across 3-4 replicate leaves from 2 plants.

The analysis of ligand-protein contacts from predicted poses pinpoints the most-contacted residues in the binding site (**Fig. S10**). Interestingly, although Y241 does not engage in direct polar interactions with the ligand, its high contact frequency in IFD suggests a strategic spatial orientation acting as a structural “pillar” (**Fig. 3C**). It appears to be part of an aromatic cluster involving F226, F239 and F212 at the center of the cavity. Interestingly, the binding pocket is flanked by three highly conserved tryptophan residues (W202, W207, and W264). This pattern of aromatic residues contributes to the geometry of the pocket, influencing its shape and driving ligand complementarity (**Fig. 3C**). Hydrophobic/aliphatic residues involved in frequent interactions include L154, L182, and I184 at the extremity of the pocket, and M260 and L274 at its center (**Fig. 3C**). A contact analysis to identify specific polar interactions (hydrogen bonds and salt bridges) involving the ligand headgroup revealed S224 and Y222 as the residues most frequently engaged by the ligand headgroup (**Fig. 3D**), suggesting they are potential anchors for the carboxylate moiety. As these residues are located in the upstream part of the flexible L2 loop (**Fig. 3C**), these findings imply that the loop plays a direct role in stabilising the bound state. In **Fig. 3B**, we report the consensus binding poses showing all interactions. In addition to those already mentioned, P225 at the upstream end of the flexible loop and H273 from the opposing β-sheet are predicted to engage with the ligand headgroup (**Fig. 3C, D**). A pair of conserved hydrophobic residues (I-I or F-F) is located at positions 215 and 218, further downstream in the loop, and may further stabilise the bound state (**Fig. 3C, D**). To experimentally evaluate the predicted interactions, we mutated single or multiple residues in *At*LORE and assessed its receptor activity upon transient expression in *N. benthamiana* by measuring 3-OH-C10:0-induced ROS production (**Fig. 3E**). Like other mutations of hydrophobic residues at the extremity or core of the pocket, the L182A or I262A mutations reduced activity, but the M260A mutation did not. Introducing a bulkier residue at G192, L182, or I262 abolishes activity, presumably by preventing the ligand tail from fully entering the pocket. Mutations in one or more residues of the aromatic cluster (Y241A, F226A, F239A, or F212A) abolish LORE activity, supporting a critical role for this structural element. Mutations W202A, W264A, and Y247A also abolish receptor function, supporting a critical structural role for these conserved aromatic residues in shaping the binding pocket. In addition to S224A, mutation of either one of the residues predicted to interact with the ligand headgroup (Y222A/H, P225A, or H273A/D) abolishes LORE receptor activation. Interestingly, mutations I215A and I218A, but not S216A, in the downstream part of the L2 loop also reduce LORE activity, albeit to different degrees, suggesting a role for these hydrophobic residues, particularly the one at position 215, in stabilising the bound complex. Additionally, we assessed the roles of loop residues D219, S221, and V223, which are highly conserved across 3-OH-C10:0-binding LORE orthologs and non-binding SD1-23 orthologs (**Fig. 2A**), suggesting a conserved role of the loop beyond ligand binding. In support of this notion, the D219A and V223G mutations strongly reduce 3-OH-C10:0-induced receptor activity, while the D219A-S221A double mutation abolishes it (**Fig. 3E**). Overall, the mutational analysis strongly supports the predicted binding mechanism: the acyl tail docks into the hydrophobic core of the binding site and interacts with the lower part of the flexible L2 loop, particularly I215 and F212, whilst the head group forms polar interactions involving the upstream part of the flexible L2 loop to stabilise the bound state. Furthermore, mwSuMD simulations and the Boltz-2-generated LORE ensemble highlighted the highly dynamic nature of the L2 loop (**Fig. S11**). Together, these data suggest that the loop has two functions in ligand binding: it acts as a dynamic gate that regulates access to the pocket, and it provides essential anchoring points once the ligand is bound.

### Ligands with 3-OH-C10:0 tails and a bulky headgroup are antagonists of LORE signalling

The proposed binding mechanism is also consistent with the observation that modifications to the carboxylate group impair the eliciting activity of 3-OH-C10:0 (Kutschera *et al*., 2019). Small modifications reduce activity, showing that these 3-OH-C10:0 analogues can still bind to the *At*LORE-ECD and activate the receptor, albeit less efficiently. By contrast, 3-OH-C10:0 analogues with complex headgroup modifications are mostly inactive, either because they cannot bind or they bind but cannot activate the receptor. To address these hypotheses experimentally, we tested whether complex headgroup 3-OH-C10:0 analogues can outcompete free 3-OH-C10:0 when applied in 1000-fold excess. First, we confirmed that complex headgroup 3-OH-C10:0 analogues do not activate LORE signalling even at the very high concentration required for this assay. Indeed, no response was observed upon application of 30 µM of the complex headgroup analogues *t*But-3-OH-C10:0, Et_2_N-3-OH-C10:0, PyrN-3-OH-C10:0, as well as decandioic acid, the non-binding control (Kutschera *et al*., 2019), in *At*LORE-overexpressing *lore*-1 seedlings (**Fig. 4A**). A subsequent dose of 30 nM free 3-OH-C10:0 triggered a response in the decandioic acid-pretreated seedlings similar to the solvent control, while no or a very low response was observed in case of the other three analogues (**Fig. 4A**). Similar results were obtained when the competitors and free 3-OH-C10:0 were applied simultaneously (**Fig. 4B**). This demonstrates that the complex headgroup 3-OH-C10:0 analogues bind to the *At*LORE-ECD but interfere with receptor activation. We then modelled the binding of the strongest agonist (3-OH-C10:0) and the three antagonists (*t*But-3-OH-C10:0, Et_2_N-3-OH-C10:0, and PyrN-3-OH-C10:0) using Boltz-2 cofolding to investigate their binding modes (**Fig. 4C**). Notably, all ligands are predicted to occupy the validated binding pocket. Across all complexes, the aliphatic tails adopt a conserved orientation, extending toward the bottom of the pocket in close proximity to residues I184 and G237 (**Fig. 2B, C**; **4C**). Moreover, the predicted binding mode is consistent with the orientation observed in the final frame of the mwSuMD R1 simulation (**Fig. 3A**). The aliphatic tail interacts closely with the critical phenylalanine triad in the core comprising residues F212, F226, and F239. While the antagonists adopt a similar orientation of the aliphatic tail, their bulky head groups are positioned at the entrance of the binding pocket, potentially affecting loop flexibility for receptor activation. In summary, 3-OH-C10:0 analogues with complex headgroup modifications bind to the receptor but fail to activate it, thus acting as antagonists of LORE signalling. Overall, our results suggest that receptor activation is a dynamic multi-step process in which ligand binding coordinates and stabilises the flexible L2 loop, thereby enabling receptor activation.

**Figure 4.**
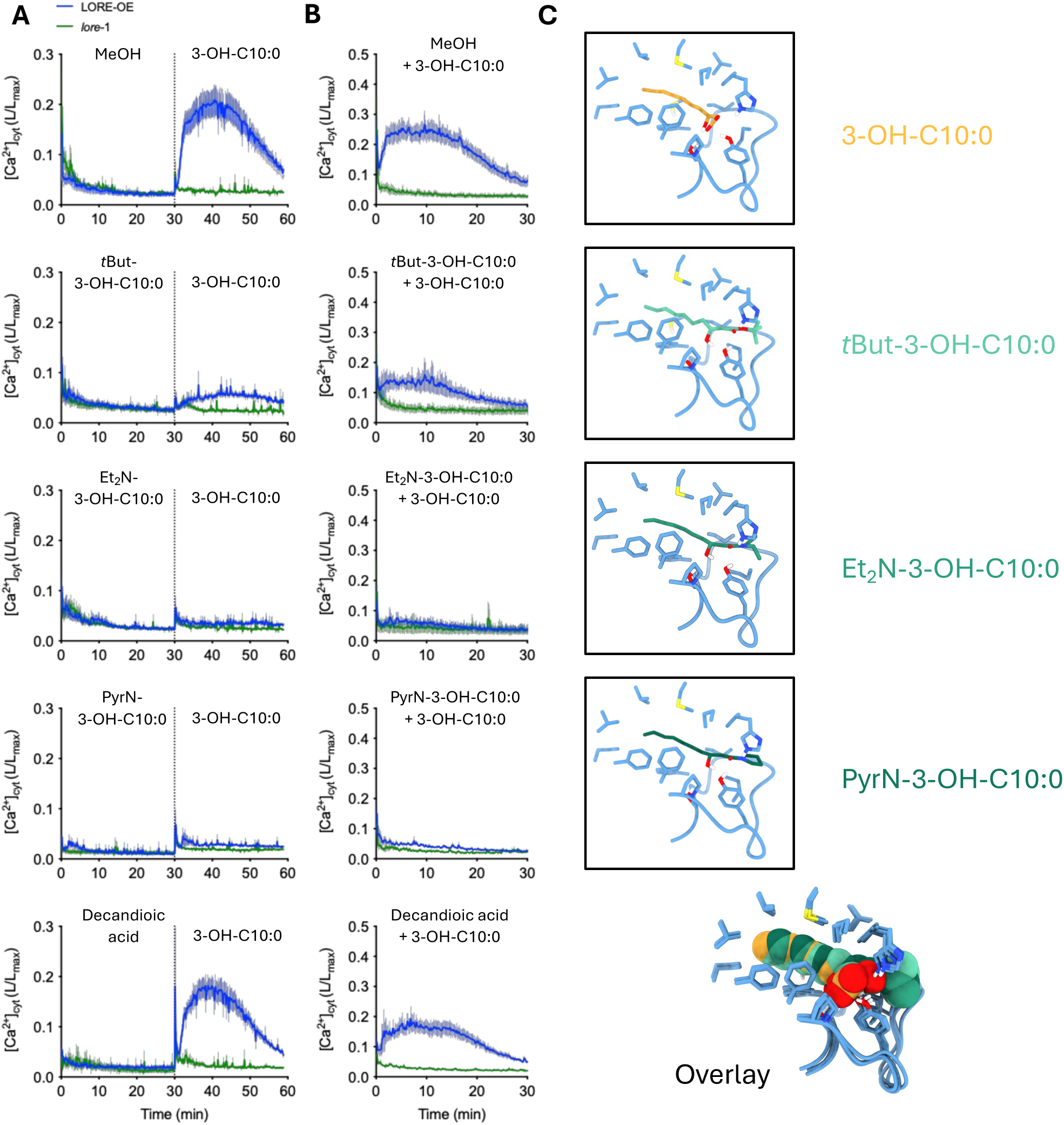
Ligands with bulky headgroups are antagonists of LORE signalling. A, **B** [Ca2+]_cyt_ kinetics were measured in LORE overexpressing (LORE-OE/*lore*-1) and *lore-*1 knockout Arabidopsis seedlings treated with 30 µM of the indicated competitors and 30 nM 3-OH-C10:0 applied sequentially (**A**) or simultaneously (**B**). Graphs show pooled data from three (**A**) or two (**B**) independent experiments, presented as mean ± SEM; total number of seedlings: LORE-OE: n≥4, *lore*-1: n ≥2. **C** Predicted binding mode of agonist and antagonist to the identified binding pocket.

## DISCUSSION

In this study, we combined structural modelling, computational ligand docking, molecular dynamics simulations, and targeted mutagenesis to characterise mc-3-OH-FA sensing by the *A. thaliana* immune receptor LORE. Our analyses identify a hydrophobic pocket in the L2 domain as the primary binding site, reveal a dynamic mechanism in which the acyl tail enters the pocket first and the carboxylate headgroup is subsequently anchored through polar interactions with the flexible L2 loop, and demonstrate that complex headgroup-modified 3-OH-C10:0 analogues act as antagonists of LORE signalling.

The sequence- and structure-based analyses of the binding pocket provide a mechanistic explanation for the inability of *At*SD1-23 to sense mc-3-OH-FAs (Shu *et al*., 2026a), despite high overall ECD sequence similarity. Several SAPs that distinguish *At*SD1-23 from *At*LORE are concentrated in the L2 binding pocket, and our mutational analyses show that individual SAPs can significantly impair ligand binding or receptor activity. However, no single substitution fully recapitulates the non-binding phenotype of *At*SD1-23; rather, it is the combined effect of multiple pocket-resident SAPs that renders the paralog insensitive to mc-3-OH-FAs. Conversely, converting all seven pocket-resident SAPs in the *At*SD1-23 L2 domain to the corresponding *At*LORE residues confers full 3-OH-C10:0 sensing capacity, whereas converting five residues confers only reduced sensing capacity, and converting a pair of critical SAPs is insufficient. This reinforces that the collective configuration of these residues, rather than any single SAP, determines binding competence. Additionally, the successful reverse engineering of the *At*SD1-23 ECD into a 3-OH-C10:0 sensing domain demonstrates that its remaining receptor features are conserved. The biological role of SD1-23, whether it is a pseudoreceptor or senses another lipidic ligand, remains to be determined.

Although definitive structural validation awaits experimental determination of the LORE-ECD structure, ideally in complex with both agonistic and antagonistic ligands, our proposed mechanism is based on convergent computational predictions validated by mutational analysis. The agreement of multiple independent computational approaches, including homology, modelling, AF2 prediction, induced-fit docking, Boltz-2 protein-ligand cofolding, and mwSuMD, together with their systematic experimental validation, lends confidence in the proposed binding mechanism.

While the proposed mechanism provides a coherent explanation for headgroup recognition and the role of the flexible L2 loop in receptor activation, it does not yet fully account for two additional features of LORE ligand specificity: the discrimination between medium-chain and long-chain 3-OH-FAs, and the strict requirement for hydroxylation at the C3 position (Kutschera *et al*., 2019). Whether these properties are governed by the dimensions of the hydrophobic pocket, by specific contacts within the cavity, or by a combination of both remains to be resolved.

The binding mechanism we describe for LORE differs conceptually from the activation paradigms established for LRR-RLKs or the SRK-SCR interaction, where ligands typically act at the receptor surface to promote co-receptor recruitment or stabilise receptor–co-receptor or receptor-dimer interfaces, respectively (Hohmann *et al*., 2017). Instead, the mc-3-OH-FA is sequestered within an enclosed hydrophobic pocket in the interior of the L2 domain, and receptor activation appears to depend on ligand-induced conformational changes within the ECD rather than on the ligand bridging an intermolecular interface. Intriguingly, intramolecular lipid sequestration as a recognition strategy has precedent in animal innate immunity. In the mammalian TLR4/MD-2 system, the acyl chains of bacterial lipid A, which include 3-hydroxy fatty acids, are buried within a deep hydrophobic cavity in the MD-2 co-receptor, and the number and length of acyl chains determine whether lipid A acts as an agonist or antagonist (Park *et al*., 2009; Ohto *et al*., 2012). By contrast, lipopeptide recognition by TLR2/TLR1 involves insertion of the acyl chains into hydrophobic pockets in both receptor subunits, such that the lipid moiety directly bridges the heterodimeric interface (Jin *et al*., 2007). LORE appears to employ the former strategy, intramolecular sequestration, consistent with the observation that mc-3-OH-FAs are small, single-chain lipids without the extended architecture needed for intermolecular bridging. Together with sphingolipid sensing by RDA2 (Kato *et al*., 2022), our findings suggest that SD-RLKs employ a structural logic for lipidic ligand recognition that is distinct from peptide perception by LRR-RLKs but shares convergent features with PRR-dependent lipid-sensing mechanisms of immune receptors in animal innate immunity.

The receptor-mediated recognition of mc-3-OH-FAs by LORE contrasts with recently described PRR-independent mechanisms by which other bacterial lipids and lipopeptides activate plant immunity. The cyclic lipopeptide surfactin from *Bacillus* spp., the diffusible signal factor (DSF) of *Xanthomonas campestris* or rhamnolipids of *Pseudomonas* spp. directly interact with the plasma membrane lipid bilayer, thereby activating mechanosensitive ion channels and immunity (Tran *et al*., 2020; Schellenberger *et al*., 2021; Gilliard *et al*., 2026; Jolivet *et al*., 2026). Typically, activation of membrane mechanosensing immunity appears to require much higher concentrations of the lipidic agonists than PRR-mediated sensing. The existence of these parallel modes of lipid-mediated immune activation in plants suggests that the chemical properties of the lipidic compounds, i.e. their amphipathicity, size, and membrane-partitioning capacity, may determine whether they are perceived through PRR-based or membrane-based mechanisms.

A central question arising from our data is how ligand binding in the L2 pocket is coupled to receptor homodimerisation and activation. LORE homodimerisation through its ECD and TMD is a prerequisite for receptor function, and LORE orthologs from *A. lyrata* and *A. halleri* illustrate that ligand binding and homodimerisation can be uncoupled (Shu *et al*., 2021; Eschrig *et al*., 2024b). Attempts to model a homodimeric LORE-ECD complex in the ligand-free state using AlphaFold Multimer have been unsuccessful. We propose that ligand-induced rearrangement of the L2 loop may generate or stabilise a dimerisation interface, thereby directly coupling ligand perception to receptor activation. This model is supported by several observations: mutations that abolish headgroup anchoring by the loop, such as S224A, Y222A, and P225A, eliminate receptor activation without necessarily preventing ligand entry into the hydrophobic pocket; the mutation of conserved loop residues D219A-S221A and V223 that do not directly contact the ligand headgroup also abolishes activation; and antagonistic analogues that can occupy the pocket fail to trigger signalling, possibly by preventing productive loop rearrangement. Testing this hypothesis will require experimental validation of receptor dimerisation, ideally by determining the ligand-bound LORE-ECD structure in a dimeric state.

The observation that 3-OH-C10:0 analogues with complex headgroup modifications occupy the binding pocket, but suppress receptor activation, raises the possibility that naturally occurring headgroup-modified 3-OH-FA derivatives could modulate LORE-mediated immunity in a biological context. Several such compounds have been identified in bacteria and shown to exhibit low or no eliciting activity in *A. thaliana* (Kutschera *et al*., 2019). Whether these or related compounds act as competitive antagonists of LORE *in planta* during bacterial colonisation remains to be determined. The structural framework described here provides a basis for evaluating such molecules and for the rational design of synthetic LORE agonists or antagonists with potential applications in plant protection.

PRR engineering is an advanced synthetic biology strategy for developing crops with broad-spectrum disease resistance. By modifying the MAMP-binding sites of PRRs, we can introduce or broaden recognition specificities and enhance immune signalling. While several LRR receptors that sense proteinaceous ligands have been successfully engineered to improve pathogen recognition, engineering PRRs for lipidic ligands remains unexplored (Li *et al*., 2025; Ngou *et al*., 2025; Yang *et al*., 2025; Zhang *et al*., 2025). The identification of the L2 domain as a ligand-binding module with defined specificity determinants opens avenues for structure-guided engineering of mc-3-OH-FA sensing capacity into crop species and for exploring whether related SD-RLKs employ analogous pocket-based sensing mechanisms for their lipidic ligands.

## MATERIALS AND METHODS

### Experimental methods

#### Plant and bacterial materials

*Arabidopsis thaliana* Col-0^AEQ^, *lore*-1, and *At*LORE-OE/*lore-*1 (p35S:*At*LORE:t35S) are cytosolic apoaequorin-expressing lines used to measure cytosolic calcium levels (Ranf *et al*., 2012; Ranf *et al*., 2015; Shu *et al*., 2021). Arabidopsis were grown in potting soil (ED73, Einheitserde, Germany) mixed with vermiculite (9:1) under short-day conditions (8 h light/16 h dark) at 22°C/20°C (day/night). *Nicotiana benthamiana* and Arabidopsis plants for seed propagation or in sterile liquid media were grown under long-day conditions (16 h light/8 h dark) at 23°C/21°C (day/night). *Escherichia coli* DH5α was used for cloning, and *Agrobacterium tumefaciens* GV3101 (pMP90) for transformation of *N. benthamiana*.

#### Elicitors for immune response assays

3-Hydroxydecanoic acid (3-OH-C10:0) was purchased from Matreya LLC, USA. Decandioic acid was purchased from Sigma-Aldrich, Germany. *t*But-3-OH-C10:0, Et_2_N-3-OH-C10:0, and PyrN-3-OH-C10:0 were synthesised previously (Kutschera *et al*., 2019). The proteinaceous elicitor flg22 (QRLSTGSRINSAKDDAAGLQIA) was synthesised by Pepmic, China.

### Multiple sequence alignment analysis of LORE and SD1-23 orthologs

The coding sequences of LORE and SD-23 orthologs were described previously (Shu *et al*., 2026a). For this study, ECDs were defined as extending from the signal peptide to approximately amino acids 410–440, encompassing conserved serine–glutamate motifs present in all LORE orthologs. The transmembrane and juxtamembrane regions were defined as extending from the serine–glutamate motifs to approximately amino acids 490–530, which include conserved lysine–leucine motifs. Intracellular domains were defined as extending from the lysine–leucine motifs to the glycine–arginine motifs adjacent to the stop codon. Protein sequences were aligned using the online MAFFT tool with default settings, as implemented in SnapGene v. 8.2.2 (Dotmatics). MSAs were converted to WebLogos using WebLogo 3 (https://weblogo.threeplusone.com).

### Molecular cloning

The *A. tumefaciens* binary vector pGGPXhc was derived from pICSL4723 and redesigned with unique T-DNA regions for Golden Gate cloning. For mCherry fusion protein overexpression, the 35S promoter, LacZ cassette, mCherry, and heat shock protein terminator (tHSP) (Nagaya *et al*., 2010) were assembled within the T-DNA, while for full-length SD-RLK expression, AtTCTPprom-LacZ-term was inserted along with a 35prom-mTurquoise-term cassette as transformation marker. To ensure compatibility with GoldenGate cloning, the CDSs of LORE, SD-23, and mutants thereof were synthesised without Esp3I sites in the coding sequences, as described (Shu *et al*., 2026a). DNA fragments encoding ECDs, TMDs, ICDs, or ECD subdomains L1, L2a, L2b, EGF and PAN, with Esp3I sites at both ends, were synthesised separately or obtained by PCR. For ECD-mCherry fusion protein expression in *N. benthamiana*, ECD fragments or the respective combinations of subdomain fragments were cloned into pGGPXhc-35Sprom-LacZ-mCherry-tHSP by Esp3I-based Golden Gate cloning with blue–white screening. Full-length receptors were obtained by assembling the respective ECD, TMD, and ICD fragments of each receptor, and chimera by assembling the respective ECD subdomains or mutants thereof with LORE-TMD-ICD, which were then inserted into pGGPXhc-AtTCTPprom-LacZ-term for expression in *N. benthamiana* using the same cloning strategy.

### Gain-of-function assay for 3-OH-FA-induced reactive oxygen species production in Nicotiana benthamiana

Full-length SD-RLKs or chimera and mutants thereof were expressed in leaves of 6- to 8-week-old *Nicotiana benthamiana* via *Agrobacterium*-mediated transformation. *A. tumefaciens* GV3101 (pMP90) carrying the expression constructs was grown on YEB agar supplemented with 50 μg/mL kanamycin, 30 μg/mL gentamicin and 100 μM acetosyringone at 28°C overnight. The bacterial culture was harvested and adjusted to an OD_600_ of 0.5 in infiltration buffer (10 mM MgSO₄, 10 mM MES, pH 5.5, 500 μM acetosyringone). The suspension was infiltrated into fully expanded *N. benthamiana* leaves using a needleless syringe. 2 days after infiltration, leaf disks (4 mm in diameter) were cut from the infiltrated leaf area and floated on 200 μL of water in a 96-well white plate overnight in the dark. 15-20 minutes before measurement, the water was replaced with 100 μL solution containing 200 μM luminol (Na salt; Roth, Germany) and 2 μg/mL horseradish peroxidase (Roth, Germany). ROS levels were measured using a Luminoskan Ascent 2.1 microplate luminometer (Thermo Scientific, USA) for 7 minutes to obtain background signals, with readings taken at 1-minute intervals. After treatment with 25 μL of 5-fold concentrated elicitor solution, the ROS burst was monitored for 60 minutes. Water served as the negative control. For each leaf disk, luminescence values were normalised to the average luminescence during the 5 minutes prior to elicitation. Average water control values, measured from separate leaf discs on the same plate, were subtracted, and total luminescence was calculated by summing the relative light units (RLUs) from 2 to 60 minutes after elicitation. RLU values were converted to % of the average response of the wild-type reference from the same leaf to enable comparisons across leaves and experiments.

### Measurement of Arabidopsis cytosolic calcium ion elevations

*A. thaliana* cytosolic apoaequorin-expressing lines were cultured in half-strength MS liquid medium (2.2 g/L MS salts [Duchefa Biochemie, Netherlands], 2.5 g/L sucrose, 0.195 g/L 2-(N-morpholino)ethanesulfonic acid [MES], pH 5.7) as described above. Seedlings were then transferred to a 96-well white plate and incubated overnight in 100 μL of water containing 10 μM coelenterazine-h (p.j.k. GmbH, Germany) in the dark. Luminescence signals were measured using a Luminoskan Ascent 2.1 microplate luminometer (Thermo Scientific, USA). Background luminescence was recorded twelve times at 10-second intervals prior to elicitation. Each seedling was treated with 25 μL of 5-fold concentrated elicitor solution, and luminescence was recorded every 10 seconds for 30 minutes after treatment. Water containing equivalent concentrations of MeOH or EtOH served as a negative control. To determine the total aequorin content of each seedling, 150 μL of discharge solution (2 M CaCl_2_, 20% ethanol) was added, and luminescence was recorded continuously for 2 min per well. Luminescence signals were normalised as L/L_max_ (luminescence counts per second/total luminescence counts) as described (Ranf *et al*., 2012), and total [Ca^2+^]_cyt_ signals were calculated by summing L/L_max_ values from 2 to 30 min after elicitation.

### Expression and harvesting of SD-RLK ECD proteins

Expression of recombinant LORE ECD proteins has been described previously (Shu *et al*., 2021). In brief, ECD-mCherry fusion proteins were expressed in leaves of 6-week-old *Nicotiana benthamiana* via *Agrobacterium*-mediated transformation. *A. tumefaciens* carrying the expression constructs was grown on LB agar supplemented with 50 μg/mL kanamycin and 100 μM acetosyringone at 28°C overnight. The bacterial culture was harvested, adjusted to an OD_600_ 0.5 in infiltration buffer (10 mM MgSO_4_, 10 mM MES, pH 5.5, 150 μM acetosyringone), and incubated at 28°C for 3 hours prior to infiltration. Cultures were mixed at equal volumes with *A. tumefaciens* carrying the silencing suppressor p19 to a final OD_600_ of 0.5. The suspension was infiltrated into fully expanded *N. benthamiana* leaves using a needleless syringe, and apoplastic proteins were harvested 3–4 days after infiltration. Detached leaves were submerged in Tris-buffered saline (TBS; 50 mM Tris base, 150 mM NaCl, pH 7.6) and vacuum-infiltrated to ∼26-inch Hg vacuum. After complete infiltration, excess surface buffer was removed, and the leaves were rolled and placed into a 30 mL syringe barrel positioned inside a 50 mL tube. Samples were centrifuged at 800 g for 20 minutes at 4°C to collect apoplastic washing fluid (AWF), which was subsequently clarified by centrifugation at 8,000 g for 20 minutes at 4°C. Prior to ligand depletion assays, the AWF was desalted using PD-10 columns (Cytiva, USA) and concentrated to the desired protein concentration using Vivaspin 20 centrifugal filters (30,000 MWCO; Sartorius, Germany). Expression of LORE ECD–mCherry fusion proteins was confirmed by SDS–PAGE and western blotting using an anti-RFP antibody (ChromoTek, Germany).

### Ligand depletion assay

The ligand depletion assay was described previously (Shu *et al*., 2021; Shu *et al*., 2026b). In brief, proteins in concentrated *N. benthamiana* AWFs were adjusted to the indicated concentrations with water. Protein samples were mixed with elicitors at a 9:1 (v:v) ratio, with elicitors prepared in water containing less than 1% MeOH, and incubated for 1 hour at 4°C on a rotator. Unbound ligands were separated from the mixture using Vivaspin 500 centrifugal filters with a 30,000 MWCO (Sartorius, Germany) and were collected in the filtrate. The ligands in the filtrates were detected by [Ca^2+^]_cyt_ measurements using LORE-OE and *lore*-1 seedlings.

### Computational methods

#### Molecular modelling of LORE ectodomain

##### Structure prediction and preparation of LORE ectodomain

We first retrieved the sequence of LORE/SD1-29 (O64782) from the UniProt database (UniProt, 2023). We predicted the 3D structure of the LORE ectodomain (ECD, residue range: 22-417) using MODELLER (Šali & Blundell, 1993) and two available templates from homologous proteins, the polymorphic S-locus receptor kinase 8 (SRK8: 6KYW) and 9 (SRK9: 5GYY). The multiple sequence alignment (MSA) was generated using the Multiple Sequence Viewer integrated in Maestro (Schrödinger Release 2021-1: Maestro, Schrödinger, LLC, New York, NY, 2021). LORE shares an overall sequence identity of 33% and 34% with SRK9 and SRK8, respectively. One thousand models were generated for each template individually, and the best models were selected based on the discrete optimised protein energy (DOPE) score (Shen & Sali, 2006) (SRK8-based: 39276, SRK9-based: -39550) and visual inspection. The AF2 model was retrieved from the AlphaFold (Jumper *et al*., 2021) database (https://alphafold.ebi.ac.uk/entry/O64782). We then superimposed the generated models onto the backbone atoms of the AF2 structure using the Protein Structure Alignment module in Maestro (Schrödinger Release 2024-3, Schrödinger, LLC, New York, NY, 2024). Lastly, the generated models were prepared using the Protein Preparation Wizard tool (Sastry *et al*., 2013) (Schrödinger Release 2021-1, Maestro, Schrödinger, LLC, New York, NY, 2021). The workflow ensured that bond orders were assigned, and missing hydrogens were added using Epik. It also generated the protonation and tautomeric states of each protein residue at pH = 7.4 ± 2.0. Then, hydrogen bonds and side-chain orientation were optimised using PROPKA (Olsson *et al*., 2011), followed by minimisation of hydrogens only with the OPLS4 Force Field.

##### LORE ECD pocket search and characterisation

Pocket detection was carried out on all three structural models using Fpocket (Le Guilloux *et al*., 2009), SiteMap (Halgren, 2009), and DeepPocket (Aggarwal *et al*., 2022) using default parameters. The predicted binding sites were compared for pocket size, location, druggability, and overall pocket scores. To assess evolutionary conservation, we performed an MSA of the LORE ECD binding pocket region (identified in L2) with orthologous sequences, as well as with proteins known not to bind 3-OH-C10:0 (**Fig. 2A**) (Shu *et al*., 2026a). The residues identified from this analysis were subsequently mapped onto the predicted binding pocket to evaluate their spatial distribution and structural context.

#### Sampling of the LORE-ECD:3-OH-C10:0 conformational space

The AF2 predicted structure of the LORE ECD was used as the starting model to explore the binding mode of 3-OH-C10:0. The Schrödinger Glide Induced Fit docking protocol (Sherman *et al*., 2006a; Sherman *et al*., 2006b) (Schrödinger Release 2021-1: Induced Fit Docking protocol; Glide, Schrödinger, LLC, New York, NY, 2021; Prime, Schrödinger, LLC, New York, NY, 2021) was applied to sample the residue conformations within the LORE ECD identified binding site (Nicoli *et al*., 2023a; Nicoli *et al*., 2023b). The docking grid was centered on the docked ligand, using the reference ligand position as the grid center. During the initial Glide (Friesner *et al*., 2004) docking step, ligand conformations were sampled within a 10 Å inner grid box, and multiple docking conformations were generated. Residues located within 5 Å of the ligand poses were then subjected to Prime refinement to allow local side-chain and backbone adjustments. The refined receptor–ligand complexes were subsequently re-docked using Glide in standard precision (SP) mode with a reduced grid box (5 Å) to optimize ligand placement in the flexible binding site.

#### Sampling the binding pathway of 3-OH-C10:0

##### MD system protocol

The initial coordinates of the MD simulations were retrieved from the prepared AF2-predicted model. The ligand 3-OH-C10:0 was placed at a distance of at least 25 Å from the nearest receptor atom. This ensures that the ligand is at a distance greater than the 9 Å cut-off for electrostatic interactions, as previously done in other ligand binding simulations (Majumder *et al*., 2023; Deganutti *et al*., 2025) The prepared protein and the ligand were solvated in a cubic box with a padding of 20 Å, using the TIP3P water model (Price & Brooks, 2004), and neutralized with a proper number of sodium and chloride ions. The final system box was 125.37Å x 125.29 Å x 125.29 Å for a total of 245350 atoms. The system was built with the Leap software implemented in AMBER (The Amber Molecular Dynamics Package, at http://ambermd.org) (Case *et al*., 2005) and AmberTools22 (Case *et al*., 2023). The protein was parametrised using the ff14SB force field (Maier *et al*., 2015). 3-OH-C10:0 topology, parameters and partial charges were retrieved using antechamber and parmcheck2 tools, assigning parameters from GAFF force field (Wang *et al*., 2004). ACEMD (Harvey *et al*., 2009) (Acellera, version 3.7.1) was used for the MD simulations with periodic boundary conditions.

The systems were initially equilibrated through a 1000 conjugate gradient step minimization to reduce clashes induced by the system preparation between protein and water atoms and then equilibrated with a 4 ns MD simulation in isothermal–isobaric conditions (NPT ensemble), employing an integration step of 2 fs. Initial restraints of 5 kcal mol^−1^ Å^−2^ were gradually reduced in a multistage procedure over the 4 ns: 2 ns for all protein atoms other than Cα atoms, 4 ns for the protein Cα atoms, and 4 ns for all the ligand atoms. The temperature was maintained at 298 K using a Langevin thermostat (Loncharich *et al*., 1992) with a low damping constant of 1 ps^−1^, and the pressure was maintained at 1.01325 atm using a Montecarlo barostat (Åqvist *et al*., 2004) The M-SHAKE algorithm (Kräutler *et al*., 2001) was used to constrain the bond lengths involving hydrogen atoms. Long-range Columbic interactions were handled using the particle mesh Ewald summation method (Essmann *et al*., 1995) with grid size rounded to the approximate integer value of cell wall dimensions. The cutoff distance for long-term interactions was set at 9.0 Å, with a switching function of 7.5 Å. A similar approach was used to equilibrate the system in our previous work on NbUGT72AY (Liao *et al*., 2025).

##### Multiple walker supervised molecular dynamics (mwSuMD) protocol

mwSuMD simulations (Deganutti *et al*., 2025) were used to study the binding of 3-OH-C10:0 at the LORE ECD. Three independent mwSuMD replicas were performed. The initial coordinates, topology, velocities and force field parameters were retrieved from the equilibration stage. Then, a series of short unbiased MD simulations, referred to as walkers, were performed using an integration time step of 4 fs. After completion of each batch, the best-performing walker was selected and propagated to generate a new set of walkers, matching the number and duration used in the preceding step. The single metric score (SMscore) score was used:

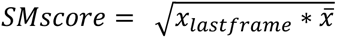

Where 𝑥_𝑙𝑎𝑠𝑡𝑓𝑟𝑎𝑚𝑒_ is the metric value in the final frame of the walker and 𝑥̅ is the average value of the same metric over the trajectory segment. In this study, the metric used to compute the SMscore was the distance between the centroid of the ligand and the centroid of residues 194, 168, 204, 243, and 264. The objective was to promote ligand binding, the metric was defined to decrease, and the walker with the lowest SMscore was propagated to the next batch.

##### MD simulations analysis

The starting AF2 model was used as the reference for structure alignment. All the simulations were analyzed with an in-house python (v. 3.12) script that exploits the following Python package of MDanalysis (Michaud-Agrawal *et al*., 2011) (v 2.4.3) for distances, RMSD calculations of protein Cα and ligand heavy atoms, root mean square fluctuation (RMSF), contact frequencies and ligand centroids during MD simulations (cutoff between any protein and ligand atoms were set to cutoff 4.5 Å).

#### Co-folding of LORE ECD in complex with different ligands

Protein–ligand complex structure prediction was performed using Boltz-2 (Passaro *et al*., 2025; Wohlwend *et al*., 2025) (v. 2.2.1). The LORE ECD sequence was provided as a single protein chain, and ligand structures were defined using SMILES strings. The ligands (3-OH-C10:0, *t*But-3-OH-C10:0, Et_2_N-3-OH-C10:0 and PyrN-3-OH-C10:0) information was retrieved from PubChem (Kim *et al*., 2025). For each ligand, co-folding simulations were carried out using the default Boltz model settings while enabling automated MSA generation via the built-in MSA server (Mirdita *et al*., 2022), and applying the Boltz potential terms for model refinement. Five models were generated per ligand, both with and without pocket constraints, to sample alternative complex conformations. Model evaluation considered predicted binding affinity together with structural confidence metrics: predicted aligned error (PAE), predicted distance error (PDE), and per-residue confidence scores (pLDDT). For the reference ligand 3-OH-C10:0, an additional set of 1250 poses was generated using Boltz-2 in unconstrained mode.

#### Consensus binding pose determination

Binding modes generated from mwSuMD simulations, IFD and Boltz-2 were jointly analysed to identify a consensus binding pose of 3-OH-C10:0 in the LORE ECD. To identify the consensus poses, we standardised atom names and file formats from the different outputs and merged all sampled poses into a single trajectory. As a reference for RMSD calculations of the ligand heavy atoms, we used the final frames from the productive replicas (R1 and R3) of mwSuMD.

#### Contact analysis

For mwSuMD trajectories, frames were retained if the ligand–pocket distance was below 6.5 Å and the ligand RMSD was below 4.5 Å relative to the reference bound pose (901 mwSuMD poses). In order to keep higher flexibility into account in our analysis, IFD poses with their ligand RMSD below 4.5 Å were analysed and compared to the final bound conformations obtained from mwSuMD replicas R1 and R3. A total of 88 IFD poses were retained for further analysis. We used the get_contacts scripts (https://getcontacts.github.io/, specifically the *‘get_dynamic_contacts.py’* and *‘get_contact_frequencies.py’*) to compute the polar interactions between 3-OH-C10:0 and the LORE ECD. The distance cutoff between any atoms of protein and 3-OH-C10:0 was set to 4.5 Å.

Visualisation of all data was done with the Matplotlib and seaborn Python library (Hunter, 2007; Waskom, 2021). Protein images were rendered with Chimera (Meng *et al*., 2023) (v1.11).

## ACKNOWLEDGEMENTS

Research of S.R. is supported by the German Research Foundation (Emmy Noether programme RA2541/1 and Collaborative Research Centre CRC924, subproject No. B10) and the Swiss National Science Foundation (grant No. 310030_208139/1). The research of A.D.P. is supported by the Leibniz Programme for Women Professors (grant: P116/2020).

---

**Figure S1.**
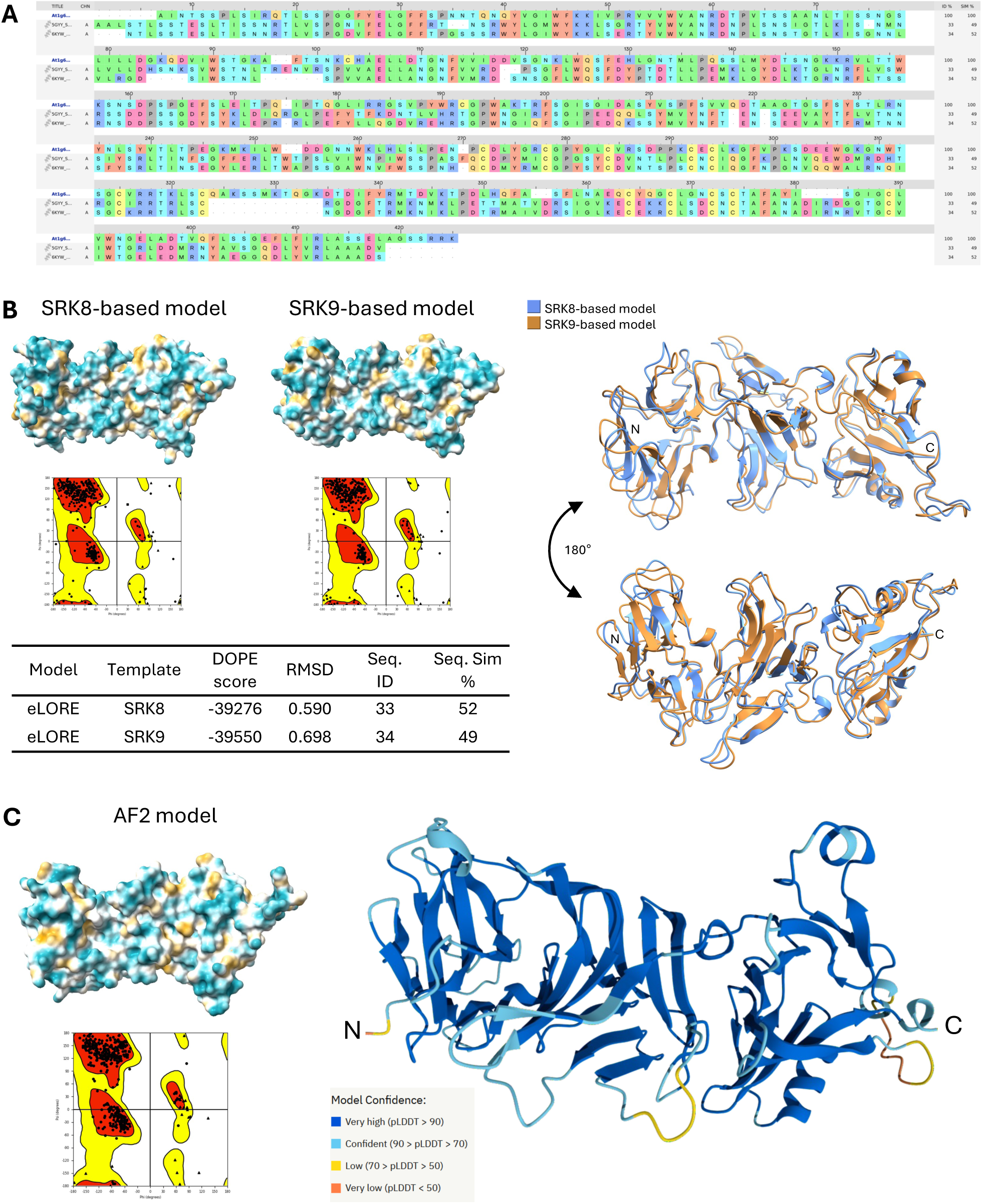
Homology-based and AlphaFold2-predicted structural models of the LORE ectodomain. **A** Multiple sequence alignment (MSA) between the ectodomains (ECDs) of LORE and SRK8 and SRK9. **B** 3D structural representation of LORE ECD homology models based on SRK8 and SRK9 shown individual in surface representation (lipohilic; left panels) and superimposed (right panel). The two models have an overall backbone RMSD of 1.83 Å and present similar DOPE score. **C** AlphaFold2 (AF2) model of the LORE ECD shown in surface representation (lipohilic; left panel) and colored according to the pLDDT confidence score (right panel).

**Figure S2.**
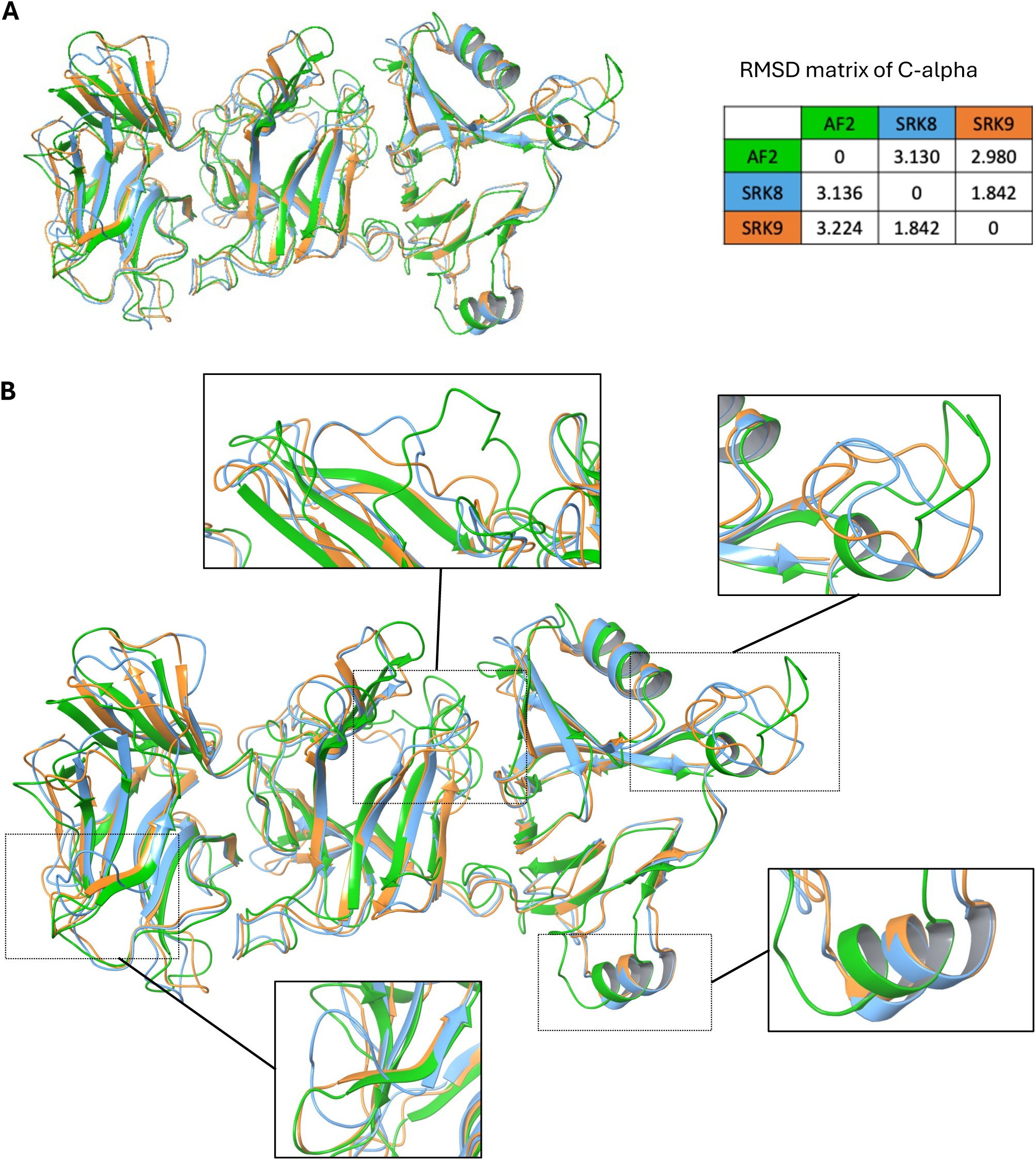
Comparison of the homology-based and AlphaFold2-predicted structural models of the LORE ectodomain. **A** Structural superposition of the LORE ECD models generated by AlphaFold2 (AF2, shown in green) and by homology modelling based on SRK8 (shown in blue) and SRK9 (shown in orange). The accompanying Cα RMSD matrix quantifies the structural similarity between the three models. **B** Enlarged views of selected regions that highlight the main local conformational differences between the models. These differences are primarily located in flexible loop regions, while the overall protein fold remains highly conserved.

**Figure S3.**
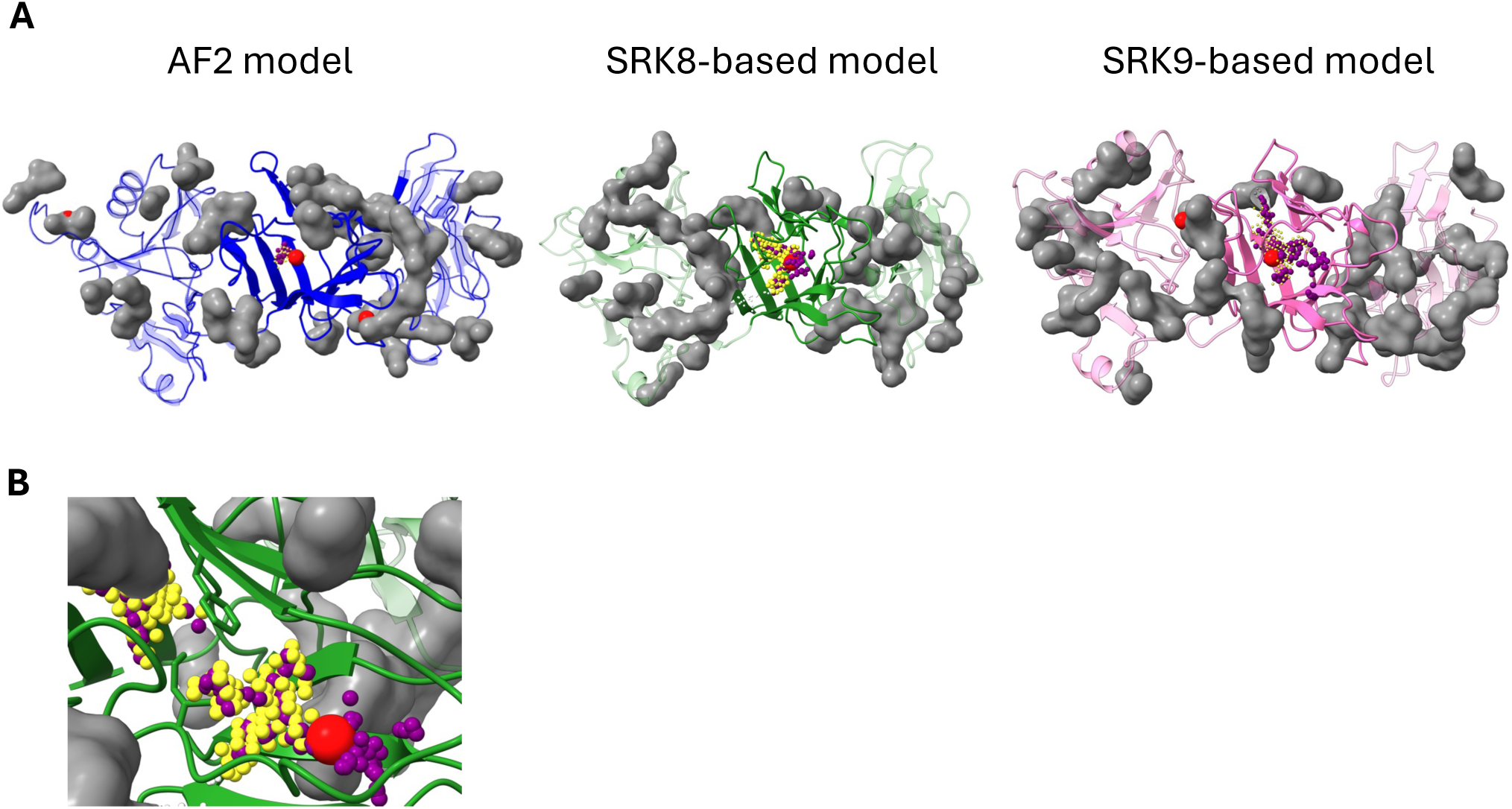
Identification and characterisation of the putative binding pocket of 3-OH-C10:0 in the LORE ECD. **A** Pocket detection was carried out on all three structural models using SiteMap, DeepPocket (red spheres), and Fpocket. Predicted sites were compared based on pocket size, location, and druggability and pocket scores, revealing a consensus binding cavity within the L2 domain. The dimensions of this pocket varied between models: the AF2-based model produced a more constricted cavity, whereas the SRK9-based model exhibited the largest pocket volume. **B** For the SRK9-based model, Sitemap and Fpocket detected two different subpockets in the primary consensus cavity because of different sidechain orientations of F231 and I215.

**Figure S4.**
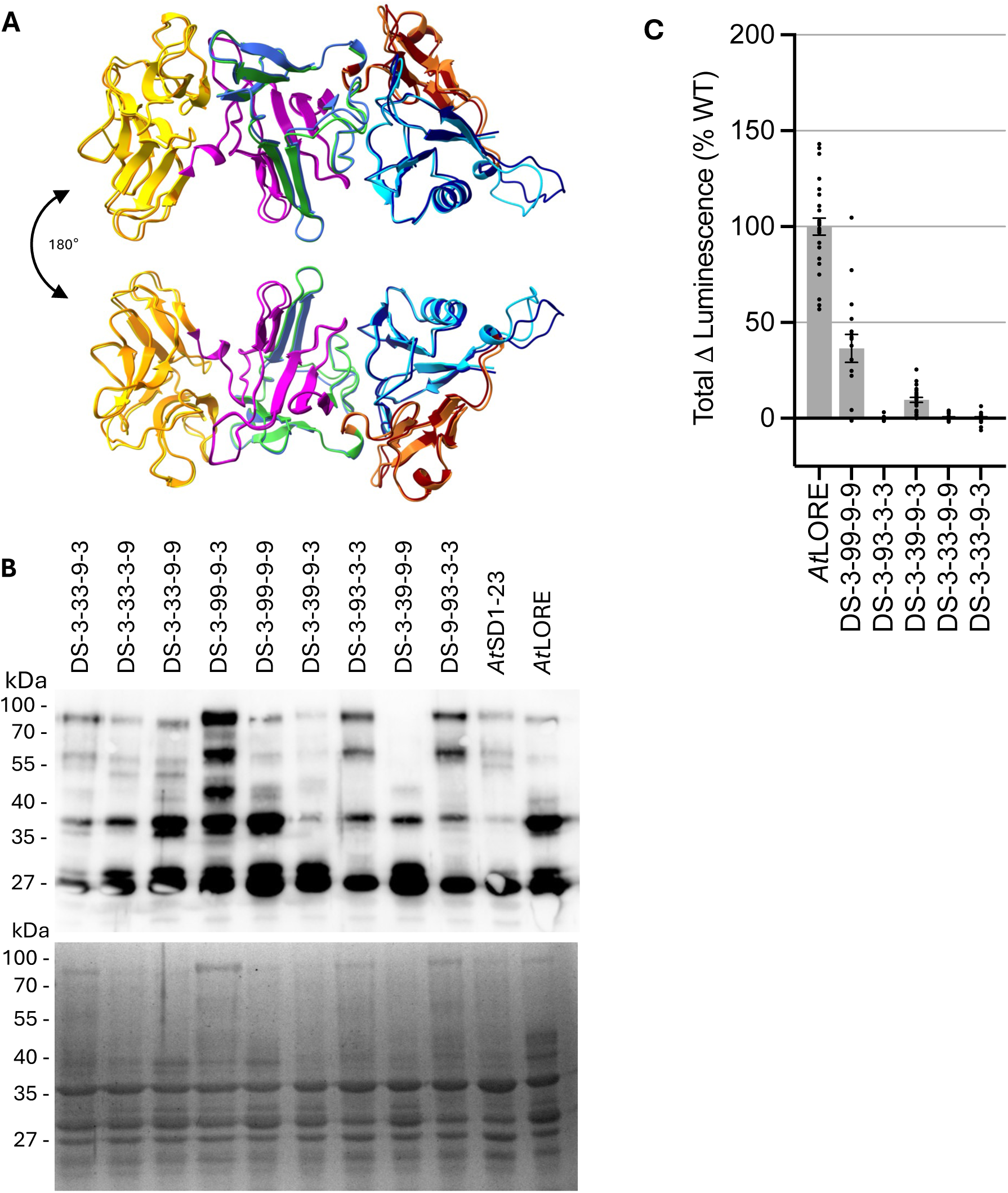
Domain swap boundaries and functional analysis of chimeric receptors in *Nicotiana benthamiana*. **A** Superimposition of AF2 models of the ECDs of *At*LORE (AF-O64782-F1-v2, 22-417) and *At*SD1-23 (AF-O64781-2-F1-v2, 42-438) with color-coded subdomains as used for domain swap experiments: L1, yellow/orange; L2a, magenta/purple; L2b, royal blue/green; EGF, turquoise/dark blue; PAN, brown/crimson red. **B** Expression of ECD-mCherry fusions with swapped subdomains in AWFs was analysed by anti-RFP immunoblot (upper panel) and Coomassie Brilliant Blue (CBB) staining of SDS-polyacrylamide gel (lower panel). AWFs were harvested 5 days after Agrobacterium infiltration in *N. benthamiana.* Desalted and concentrated AWFs corresponding to 7.5 !g total protein were loaded. The calculated molecular weights of SD-RLK ECD-mCherry fusions are 70-80 kDa. Equal sample loading was confirmed by CBB staining of the SDS-polyacrylamide gel prior to blotting. One representative immunoblot of two experiments is shown. **C** ROS production was measured 3 dpi in leaf disks from *N. benthamiana* leaves expressing *At*LORE or the indicated *At*LORE/*At*SD1-23 chimera after elicitation with 5 µM 3-OH-C10:0 or water as control. Graph shows pooled data from two independent experiments, expressed as % of wild-type response and presented as mean ± SEM with individual data points, from n = 8 leaf disks per construct across a total of 4 replicate leaves from 2 plants. DS; domain swap; L, lectin; EGF, epidermal growth factor-like; PAN, plasminogen-apple-nematode; 3 indicates subdomains of *At*SD1-23; 9 indicates subdomains of *AtLORE*.

**Figure S5.**
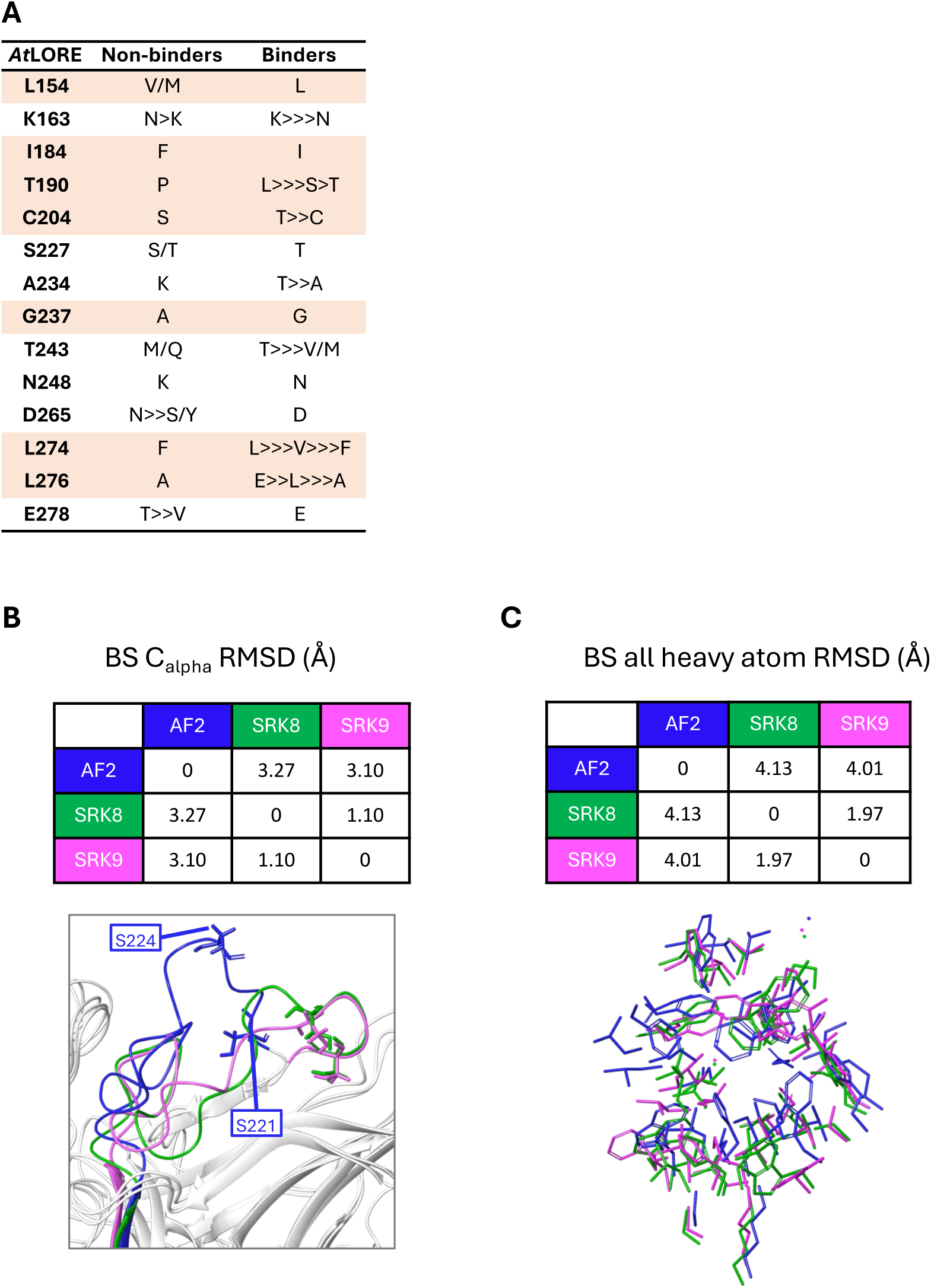
Identification of candidate residues contributing to ligand sensing in the predicted L2 binding cavity. **A** Summary of single amino acid polymorphisms located in the L2 domain of LORE-like 3-OH-C10:0 binders and SD1-23-like non-binders. Residues oriented towards the binding pocket are highlighted in orange. **B** Cα RMSD matrix and structural superposition of the L2 loop. **C** All-heavy-atom RMSD matrix and superposition of the binding site residues shown in stick representation.

**Figure S6.**
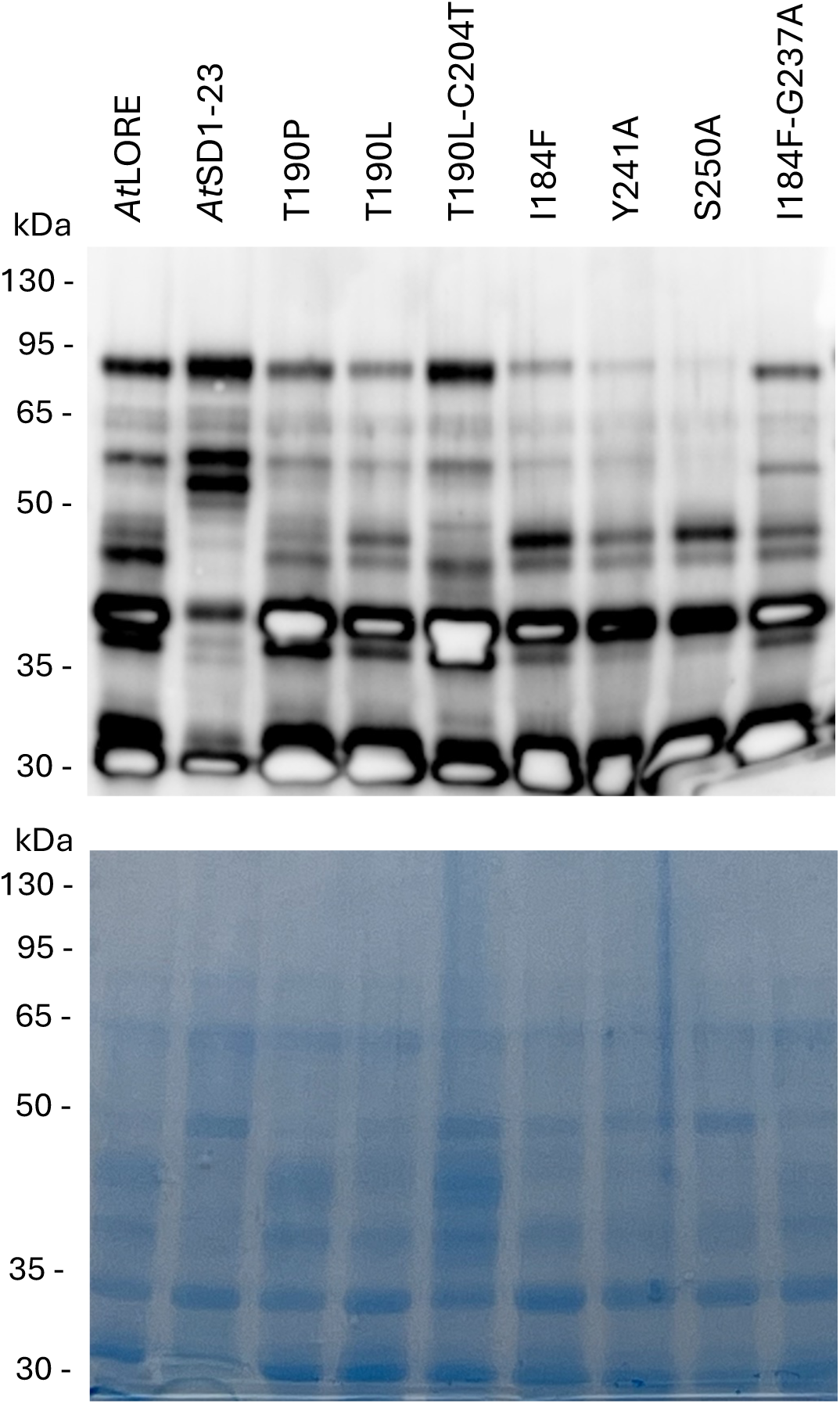
Expression analysis of mutated LORE ectodomains in *Nicotiana benthamiana*. Expression of ECD-mCherry fusions of *At*LORE, *At*SD1-23 and *At*LORE with the indicated mutations in AWFs was analysed by anti-RFP immunoblot (upper panel) and Coomassie Brilliant Blue (CBB) staining of SDS-polyacrylamide gel (lower panel). AWFs were harvested 5 days after Agrobacterium infiltration in *N. benthamiana.* Desalted and concentrated AWFs corresponding to 7.5 𝞵g total protein were loaded. The calculated molecular weights of SD-RLK ECD-mCherry fusions are 70-80 kDa. The loading of the samples was confirmed by CBB staining of the SDS-polyacrylamide gel prior to blotting. One representative immunoblot of two experiments is shown.

**Figure S7.**
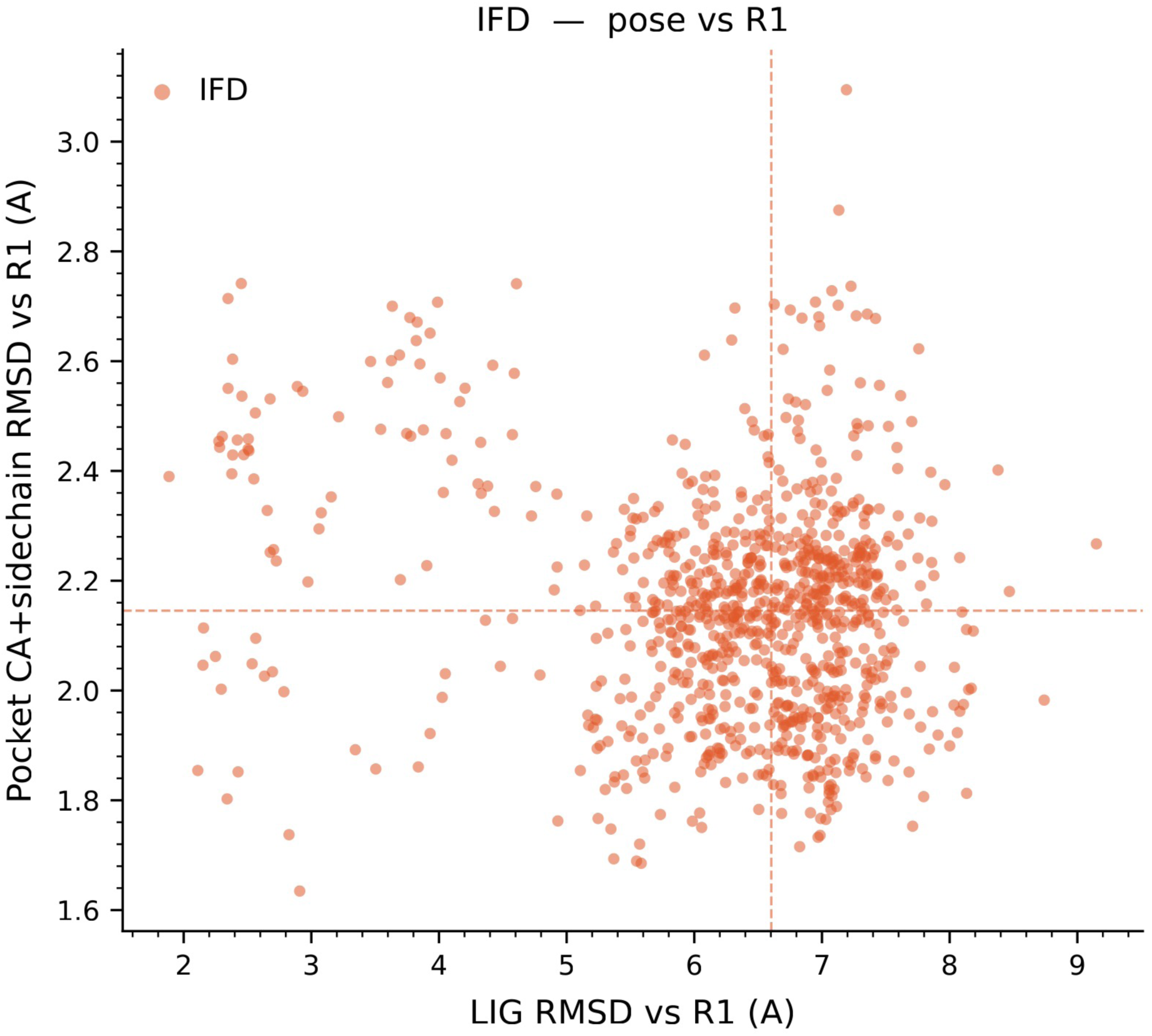
Conformational space sampled by IFD. Distribution of RMSD values for ligand heavy atoms and protein side chains across the generated poses. The last frame of R1 was used as a
reference.

**Figure S8.**
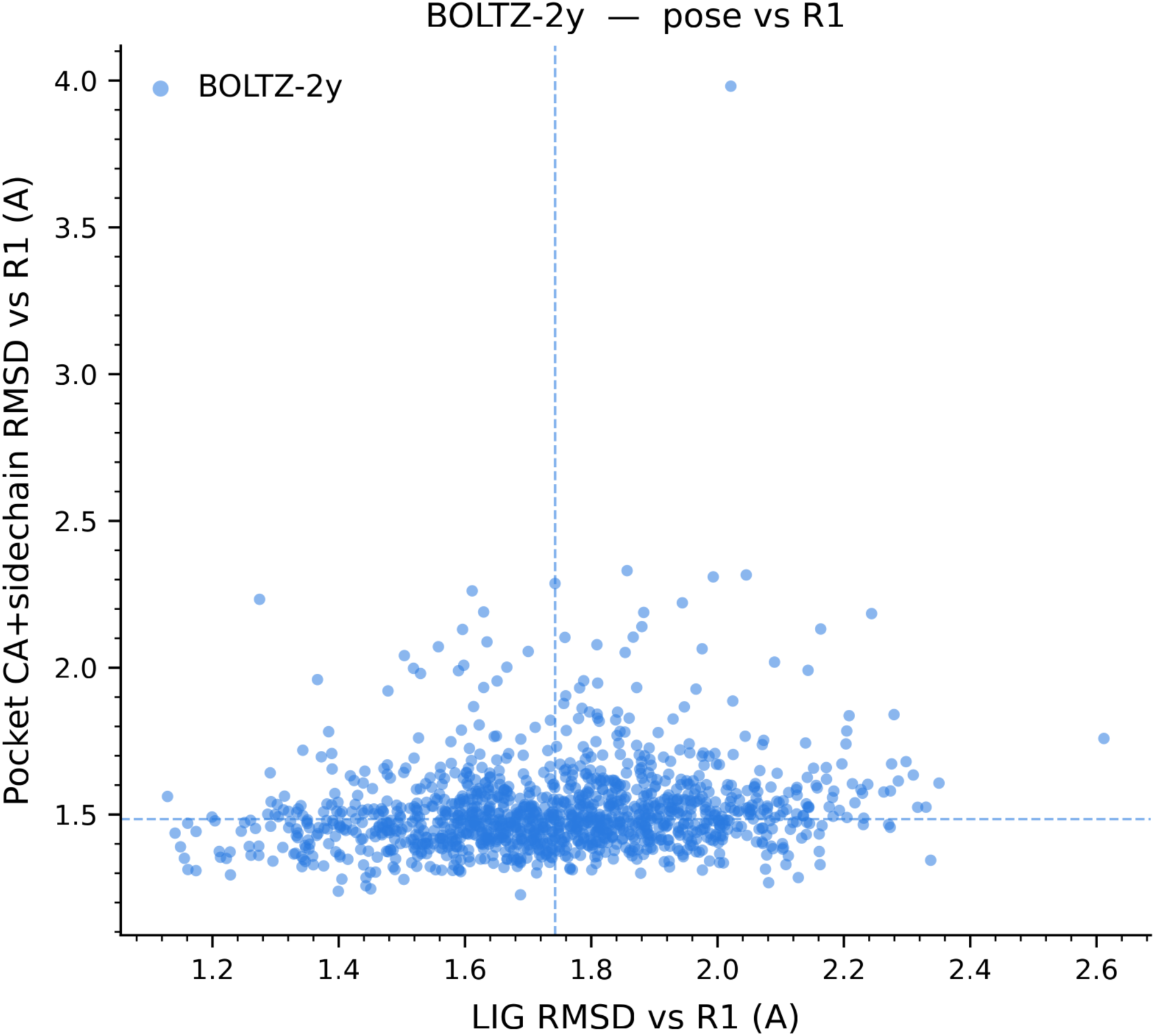
Conformational space sampled by Boltz-2. Distribution of RMSD values for ligand heavy atoms and protein side chains across the generated poses. The last frame of R1 was used as a reference.

**Figure S9.**
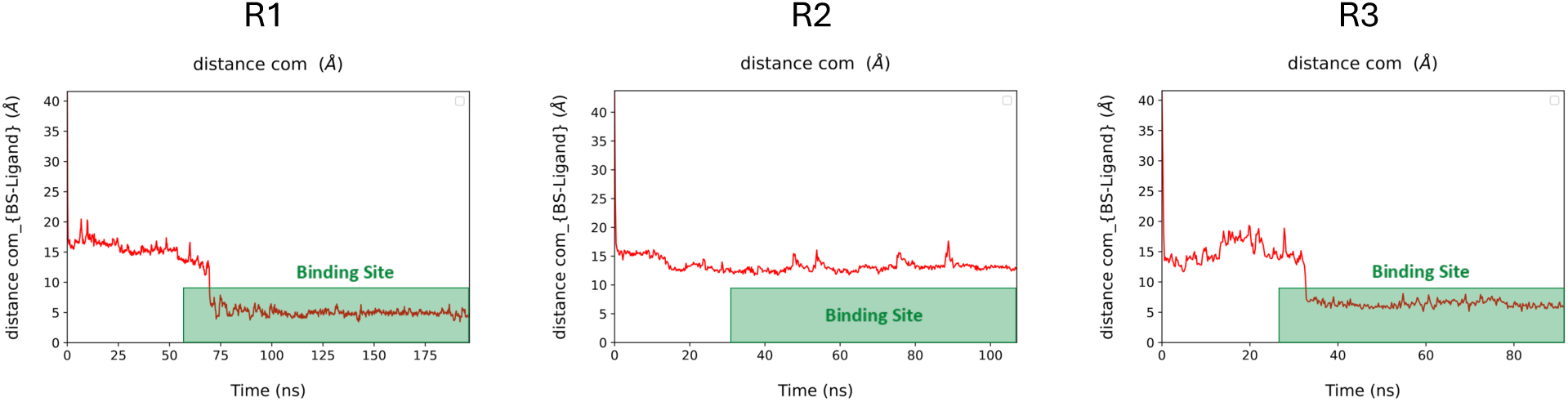
Multiple walker Supervised Molecular Dynamics simulations. Ligand RMSD during mwSuMD simulations performed in three independent replicas R1 to R3. The shaded green regions indicate frames corresponding to stable binding events within the binding site.

**Figure S10.**
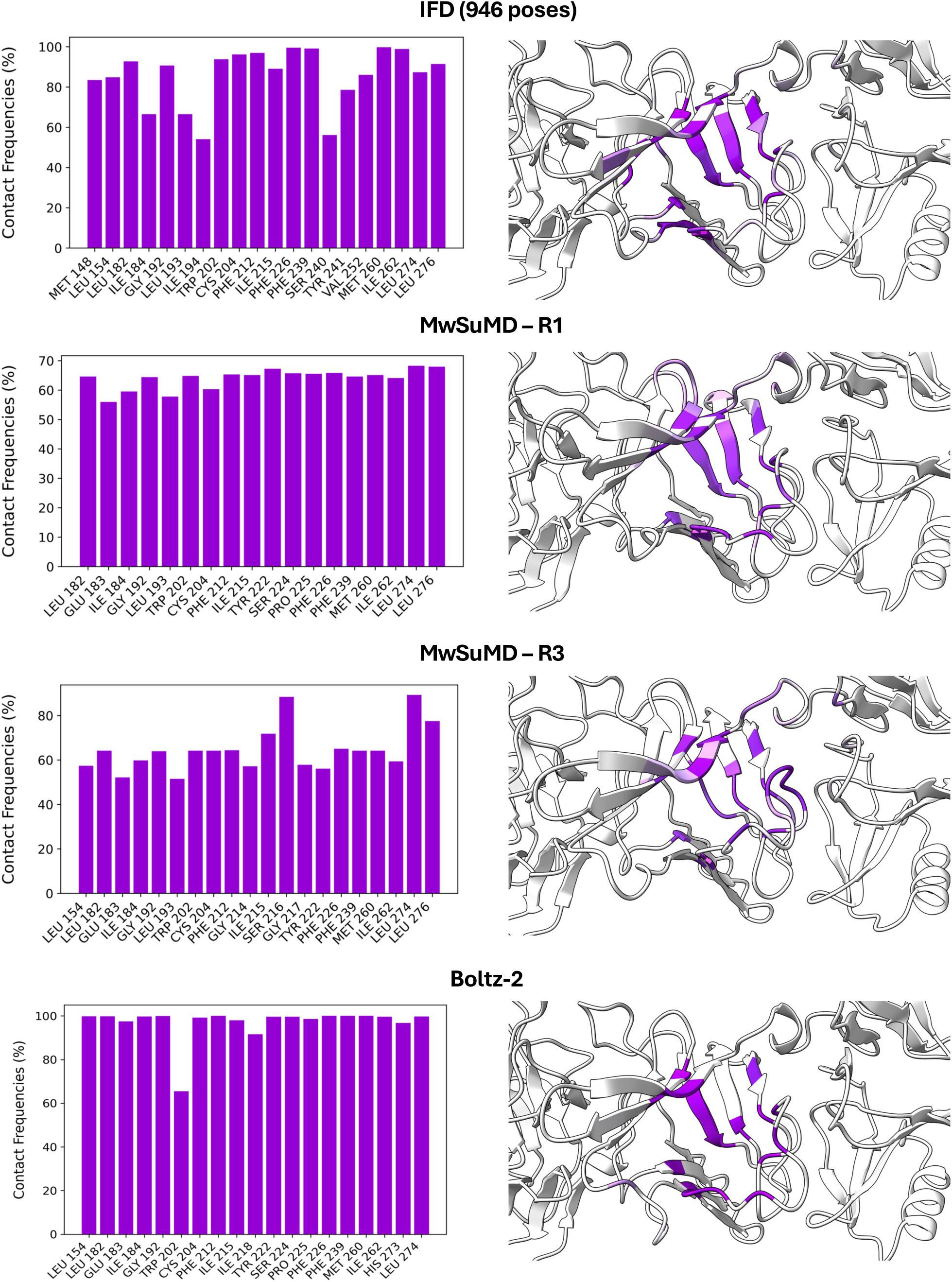
Per-residue ligand contact frequencies. Bar plots (left) show the contact frequency (%) for each LORE receptor residue within the distance cutoff, computed across all poses for IFD (top row), mwSuMD R1 (second row), mwSuMD R3 (third row), and the Boltz-2 sampling set (bottom row). Only residues with high contact frequencies are shown. The corresponding structural representations (right) highlight the contacting residues mapped onto the LORE receptor, colored in purple, with contact frequency encoded by color intensity. The LORE receptor is shown as grey cartoon.

**Figure S11.**
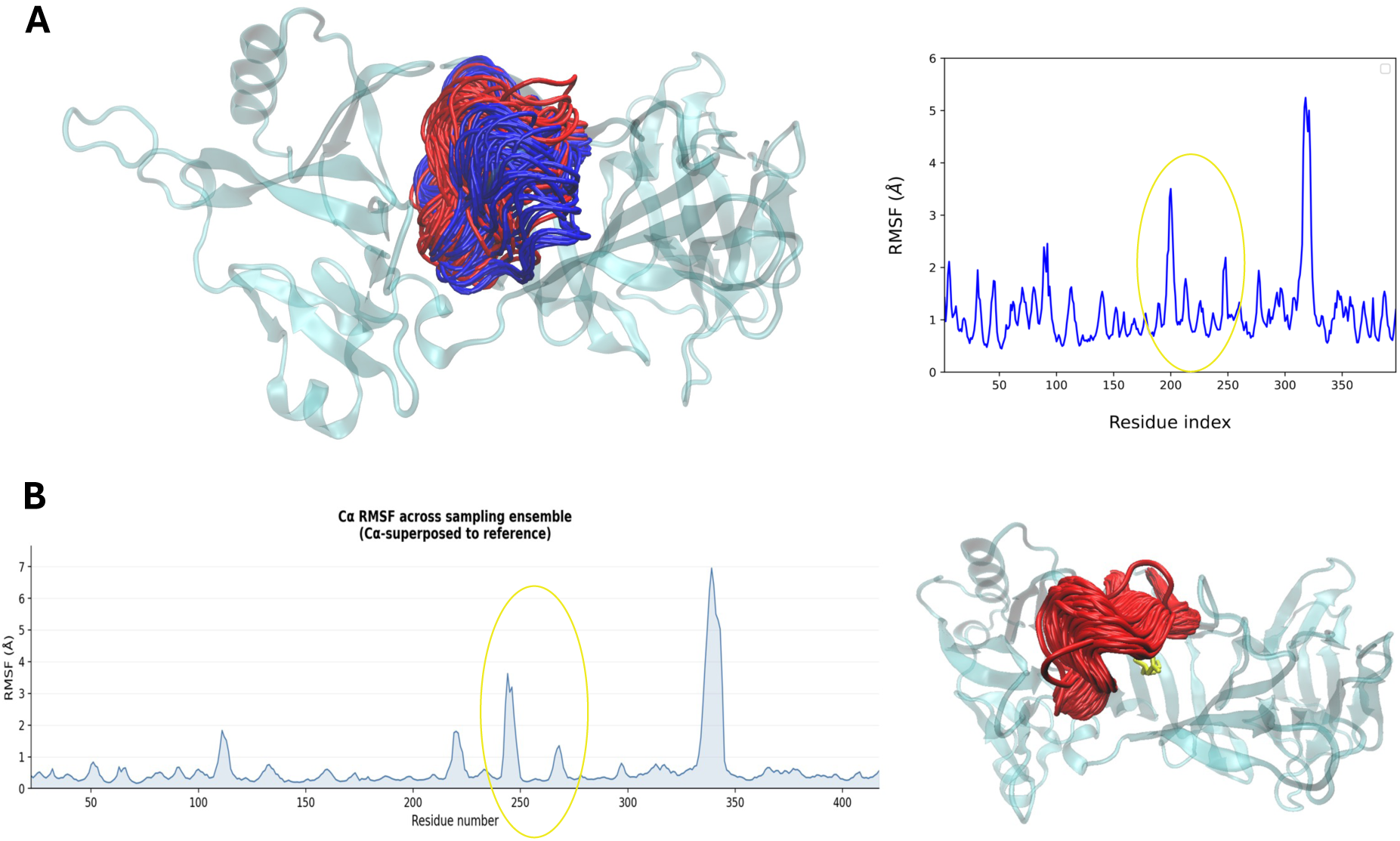
Flexibility of the L2 loop. **A** The loop region surrounding the binding site displayed significant conformational variability during the mwSuMD simulation. **B** RMSF profile and superimpositions of 1250 conformations predicted by Boltz-2. Similar to the mwSuMD, the L2 loop shows high flexibility.

